# Neurophysiological mechanisms underlying post-stroke deficits in contralesional perceptual processing

**DOI:** 10.1101/2023.12.12.571233

**Authors:** Daniel J. Pearce, Ger M. Loughnane, Trevor T.-J. Chong, Nele Demeyere, Jason B. Mattingley, Margaret J. Moore, Peter W. New, Redmond G. O’Connell, Megan H. O’Neill, Dragan Rangelov, Renerus J. Stolwyk, Sam S. Webb, Shou-Han Zhou, Méadhbh B. Brosnan, Mark A. Bellgrove

## Abstract

Slowed responding to sensory inputs presented in contralesional space is pervasive following unilateral cerebral stroke, but the causal neurophysiological pathway by which this occurs remains unclear. To this end, here we leverage a perceptual decision-making framework to disambiguate information processing stages between sensation and action in 30 unilateral stroke patients (18 right hemisphere, 12 left hemisphere) and 27 neurologically healthy adults. By recording neural activity using electroencephalography (EEG) during task performance, we show that the relationship between strokes in either hemisphere and slowed contralesional response times is sequentially mediated by weaker target selection signals in the contralateral hemisphere (the N2c ERP), and subsequently delayed evidence accumulation signals (the centroparietal positivity). Notably, asymmetries in CPP and response times across hemispheres are associated with everyday functioning. Together, these data suggest a plausible neurophysiological pathway by which post-stroke contralesional slowing arises and highlight the utility of neurophysiological assessments for tracking clinically relevant behaviour.

---

Stroke is the third-leading cause of combined death and disability in adults^1^, and is associated with both physical and cognitive impairment. One of the most reliably reported cognitive phenomena following unilateral hemispheric stroke is impaired processing of sensory inputs. This impairment typically manifests in slowed responses to sensory stimuli presented to contralesional space, and is independent of response modality, stroke type, time since stroke, and the presence of overt perceptual deficits (e.g.,^2–9^). Such impairment compromises everyday living activities, including self-care and driving^10,11^, and may hinder higher cognitive functioning^12,13^. A critical bottleneck in advancing post-stroke care is a lack of clarity regarding precisely which neural processes result in slowed responses. Indeed, response speed represents the culmination of a complex series of attentional, sensory, decisional, and motor processes^14–21^. Thus, defects arising at any of these stages could potentially drive superficially homologous impairments in overall speed.

Previous work with EEG has revealed differences in sensory processing between stroke patients and healthy controls. Most consistently, event-related potentials (ERPs) related to early perceptual (e.g., the N100)^22–26^ and decisional processes (e.g., the P300 and more specifically the P3b subcomponent)^24,25,27–32^ are suppressed or delayed in the lesioned hemisphere while stroke patients undertake speeded tasks, relative to ERPs in the healthy hemisphere. However, the majority of studies have selectively examined patients with right hemisphere lesions in the context of spatial neglect^23,25,26,28,31,33–39^, which does not wholly account for the broader incidence of slowed contralesional responding. Further, these stroke-related ERP changes are not typically associated with concurrent behavioural performance (though some relate to non-contemporaneous pen-and-paper measures, see^26,27,33^), or are only in right hemisphere stroke under very specific behavioural conditions (i.e., for left space after an invalid pre-cue^34^). Establishing these relationships may be possible with the use of paradigms that do not feature sudden changes in stimulus intensity, thereby improving the signal-to-noise ratio of overlapping non-sensory-evoked signals by reducing transient, purely sensory electrophysiological activity at the scalp^40^.

To this end, our group has developed a bilateral variant of the random dot motion task used to probe perceptual decision-making in humans and animal models (e.g.,^41^). The paradigm employs a gradual progression to target motion, which reduces the interference from sensory-evoked potentials. In the context of EEG, this paradigm can then separate neural responses related to early target selection (N2c and N2i^19,20^) and evidence accumulation (centro-parietal positivity, CPP^18,19,42,43^). Each of these metrics has been shown to independently predict response times, and they present advantages over previous metrics such as the N2pc (e.g.,^19^) and the P300 (e.g.,^44,45^). For example, the N2c is a hemisphere-specific early target selection signal that modulates later evidence accumulation and does not rely on the calculation of difference waves, providing greater sensitivity and thus explanatory power for behaviour^19^. Similarly, the CPP is a sensitive metric of supramodal decision formation which shares key features with hypothetical decision variables predicted by sequential sampling models^45,46^ and with invasive electrophysiological signals of evidence accumulation recorded in monkeys^42,43^. Further, there is now considerable evidence supporting the CPP’s causal association with response speed and interindividual differences in both younger^18,43,47–55^ and older adults^20,56^. Thus, this paradigm may enable the establishment of a neurophysiological pathway between unilateral brain lesions and slowed response speeds to contralesional stimuli.

Here, by recording neural activity using electroencephalography (EEG) during task performance, we use the bilateral random dot motion task to show that contralesional slowing in response time (RT), irrespective of the lesioned hemisphere, is causally related to a disruption in the dynamics of early target selection signals, which consequently perturbs evidence accumulation processes. Crucially, the disrupted evidence accumulation underpinning contralesional slowing correlates with self-reported everyday functioning in stroke survivors. These findings identify the causal neurophysiological mechanisms giving rise to post-stroke contralesional slowing and highlight the potential utility of individualised EEG-based measures in stroke assessment and rehabilitation.

## Results

### Unilateral Stroke Compromises the Speed of Contralesional Responding

We implemented a bilateral variant of the widely used random dot motion task^18,19^ (see Fig. 1), while brain activity was continuously recorded using high-density scalp EEG, and while real-time eye-tracking enforced central fixation. Fifty-seven participants across three groups completed the task: 30 adults who had experienced a unilateral stroke (*n*=18 right hemisphere, *n*=12 left hemisphere; hereafter ‘patients’), and 27 neurologically healthy controls of comparable age (hereafter ‘controls’). Participants did not have clinical extinction^57^ or visual field defects that would affect their capacity to respond, and hit rate was in general at ceiling across participants (although four right hemisphere patients had a hit rate lower than 90% for left hemifield targets). For each participant, we derived single-trial metrics of response time (RT), early target selection (N2c), and evidence accumulation (CPP) to explore how these metrics differed within each hemifield as a function of stroke hemisphere.

**Figure 1.**
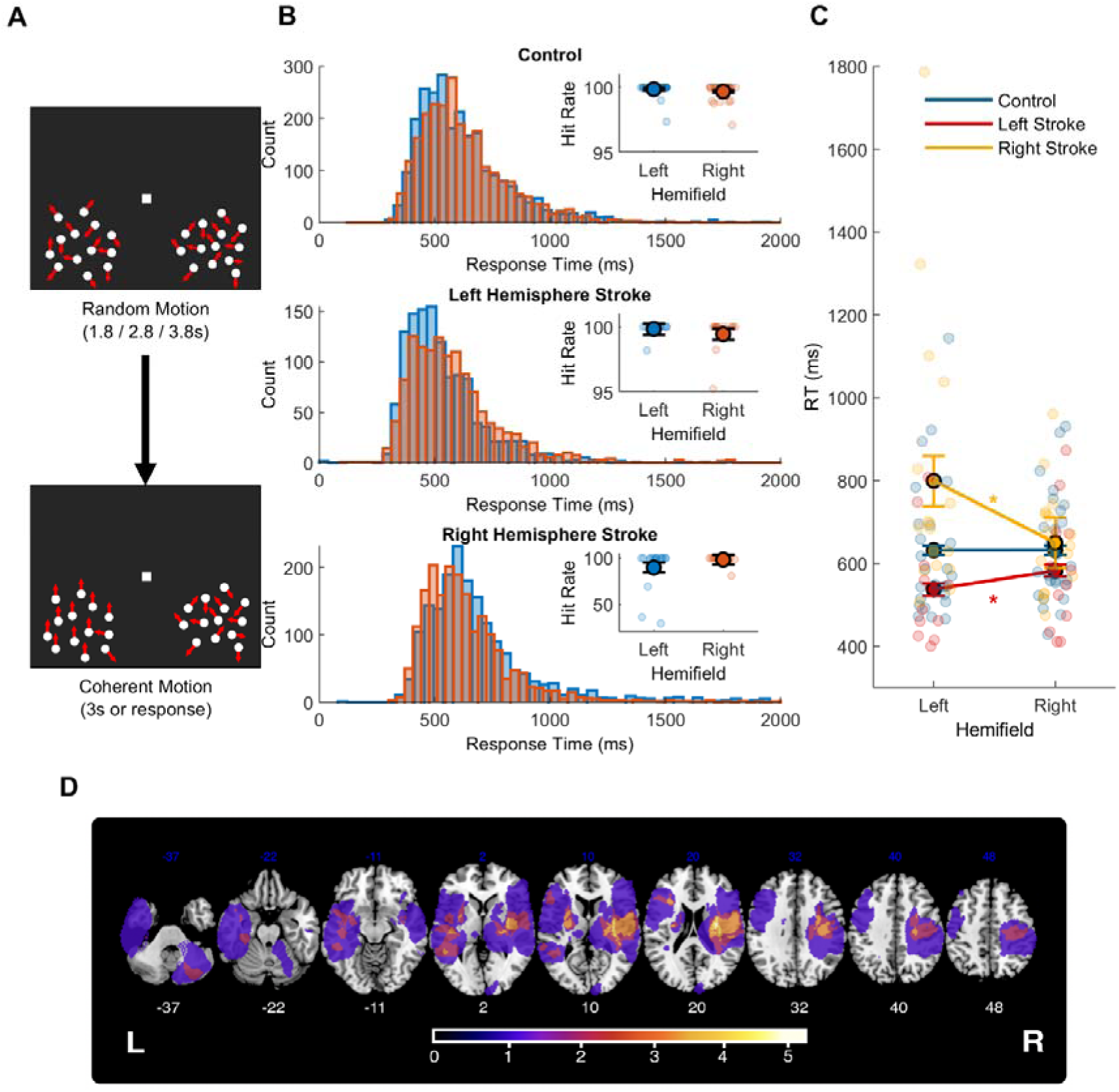
Perceptual Decision-Making Task Reveals Slowed Contralesional Responding Post-Stroke. **A.** Schematic of the BRDM task. Beginning with central fixation, each dot moved randomly at a rate of 5 degrees per second. Following a period of random motion lasting either 1.8, 2.8, or 3.8 seconds, 90% of the dots in one patch began moving uniformly, either up or down. Participants were 1 to respond to these coherent motion targets with a single mouse click with their right index finger (left index finger for *n*=2 patients with right-sided hemiplegia). **B.** Performance on the BRDM task, with trials in which target motion was presented in the left or right hemifield plotted separately. Histograms depict the response time for each valid trial (i.e., where central fixation was maintained, there were no artefacts in the EEG/eye-tracking data, and a response was made between 150ms post-stimulus onset and stimulus offset) across all participants within a given group. Note that responses up to 3000ms were permitted, but for clarity only responses up to 2000ms are visualised here. Mean hit rate for each participant is depicted via inset scatterplots. **C.** Mean reaction time for left and right hemifield target trials for each group, with individual participant means plotted as single points. **D.** Lesions overlaid for n=21 patients with available clinical neuroimaging (n=11 left hemisphere, n=10 right hemisphere). Voxel colour indicates the number of overlapping lesions present at each region (range = 1-4). Lesions are coloured according to the number of overlapping voxels between participants. Individual lesion masks, network disconnection summaries, and cortical integrity statistics can be found in Supplementary Figs. 1 and 2, and Supplementary Table 3, respectively. Significant (*p*<.05) within-group differences are denoted by asterisks. Within-subject error bars are depicted.

We first utilised a linear mixed-effects model (LMM) to examine group and target hemifield differences in RT. RT was measured as the time (in milliseconds) from the onset of target motion to a response. Trials were only included if there was no ocular deviation from central fixation and there were no artefacts in the EEG/eye-tracking data throughout the trial. The model included Group (healthy control, left hemisphere stroke and right hemisphere patients), Target Hemifield (left and right), and a Group x Target Hemifield interaction as fixed effects, with a random effect to account for within-subject effects. Note that all post-hoc comparisons referred to throughout were Bonferroni-Holm adjusted.

The LMM revealed a significant main effect of Group (*F*(2,11275)=3.87, *p*=.02; see Supplementary Table 1 for detailed information for all models), but no significant main effect of Target Hemifield (*F*(1,11275)=2.27, *p*=.13). Critically, there was a significant interaction between Group and Target Hemifield (*F*(2,11275)=7.21, *p*<.001). The interaction effect was driven by the two stroke groups, who showed significantly slowed responding to contralesional, relative to ipsilesional, stimuli. On average, the right hemisphere patients responded 137 ms more slowly to left compared with right hemifield targets (left: *M*=794ms, *SD*=327; right: *M*=657ms, *SD*=120; *t*(3769)=-2.38, *p*_cor r_=.03, model estimate difference=-136, 95% CI [-248.41 −24.05]), whereas the left hemisphere patients responded 44 ms more slowly to right compared with left hemifield stimuli (left: *M*=538ms, *SD*=129; right: *M*=582ms, *SD*=147; *t*(2514)=3.43, *p*corr=.01, model estimate difference=44.76, 95% CI [16.36 73.16]). As expected, there was no significant asymmetry in response times for the controls (left: *M*=632ms, *SD*=165; right: *M*=632ms, *SD*=130; *t*(4992)=0.07, *p*_cor r_=.94, model estimate difference=0.73, 95% CI [-19.57, 21.04]). An exploratory analysis suggested that such contralesional slowing was seen in both those with and without neglect (see Supplementary Results 1).

Overall, these results indicate that stroke – irrespective of the hemisphere of damage, or the presence or absence of a neglect – compromises the speed of response to contralesional stimuli, and that the BRDM task can detect these slowed responses. These results are shown in Fig. 1.

Next we sought to understand the sequence of neural events that give rise to the contralesional slowing observed in both stroke patient groups. Based on our past work, we hypothesised that contralesional slowing of response times could be influenced by variation in the dynamics of the N2, CPP, or both, as a function of target hemifield and lesioned hemisphere.

### Early Contralateral Target Selection Signals Have Lower Amplitudes in the Lesioned, Relative to Non-Lesioned, Hemisphere

We extracted target selection signals at posterior electrodes contralateral (N2c) and ipsilateral (N2i) to the target hemifield, as per our previous work^18–20,44^. Notably, our method allows extraction of hemisphere-specific N2 signals which can be used to compare target selection as a function of hemifield, and demonstrates robust relationships to response time.

#### N2c amplitude

To examine group and target hemifield differences in N2c magnitude we computed an LMM based on single-trial estimates of peak amplitude. As with RT, the LMM showed a significant Group x Target Hemifield interaction (*F*(2,8599)=6.25, *p*<.01; Fig. 2A; Supplementary Table 1), whereby both patient groups showed reduced amplitudes for contralesional, compared with ipsilesional, stimuli (i.e., attenuated signal in the lesioned hemisphere; Supplementary Table 1). The right hemisphere patients had 9.96 µV/m^2^ greater N2c amplitude for right, compared with left, hemifield targets (left: *M*=-8.39, *SD*=8.99; right: *M*=-18.34, *SD*=13.75; *t*(3131)=2.73, *p*_co r r_=.027, model estimate difference=-10.02, 95% CI [-17.21,-2.83]), whereas the left hemisphere patients showed an 7.61µV/m^2^ greater (i.e. more negative) N2c amplitude for left relative to right hemifield targets (left: *M*=-19.17, *SD*=8.28; right: *M*=-11.56, *SD*=9.76; *t*(1800)=-3.48, *p*_cor r_=.01, model estimate difference=8.31, 95% CI [3.08, 13.54]). N2c amplitude was equivalent for left and right hemifield targets in controls (left: *M*=-19.21, *SD*=13.20; right: *M*=-17.28, *SD*=15.44; *t*(3668)=-0.65, *p*_co rr_ =.52, 95% CI [-4.41, 8.48]). Finally, there was no significant main effect of Group (*F*(2,8599)=1.35, *p*=.26) or Target Hemifield (*F*(1,8599)=0.005, *p*=.99).

**Figure 2.**
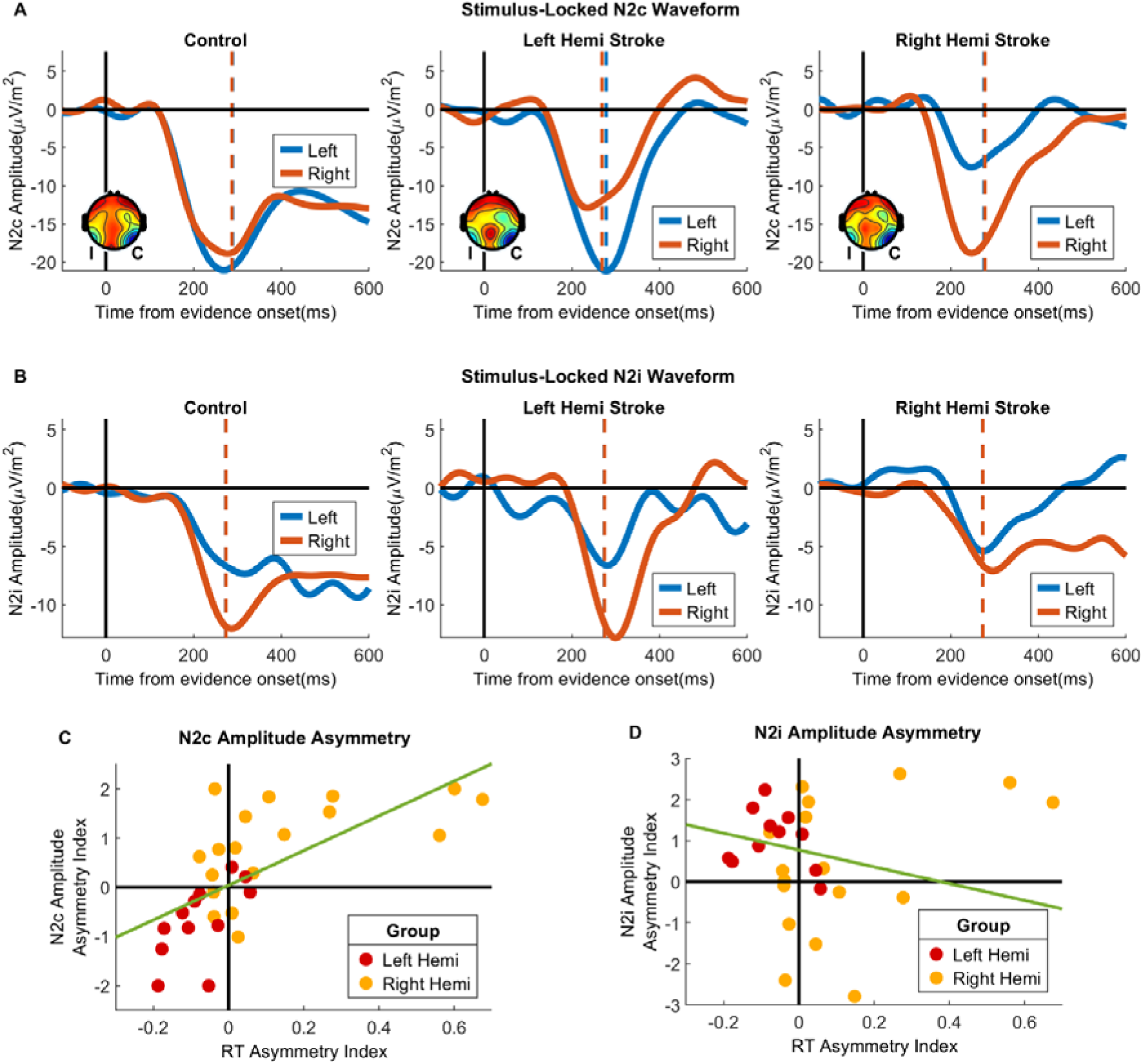
Reduced N2c, but not N2i, Amplitude in the Stroke-Affected Hemisphere Indexes Slowed Responding to Contralesional Space. **A.** N2c waveforms extracted from temporal electrodes contralateral to target hemifield (T7 and T8 for right and left hemifield targets, respectively). Inset topoplots show average scalp electrical activity during 250-280ms windows post-stimulus onset for each group separately. Note that ‘I’ and ‘C’ denote ipsilateral and contralateral signals (such that ipsilateral shows the grand average of left hemisphere activity for left hemifield targets, and right hemisphere activity for right hemifield targets, and vice versa). **B.** N2i waveforms extracted from temporal electrodes ipsilateral to target hemifield (T8 and T7 for right and left hemifield targets, respectively). **C, D.** The relationship between RT asymmetry index and N2c amplitude asymmetry index for patients. A least squares regression line is fitted for reference.

#### N2c peak latency

We then used a two-way mixed-effects ANOVA on participant-level means to examine the effects of Group and Target Hemifield on the peak latency of the N2c (i.e., the time from stimulus onset until the peak amplitude of the average waveform). There were no significant main effects of Group (*F*(2,54)=0.50, *p*=.61) or Target Hemifield (*F*(1,54)=0.08, *p*=.78), and no significant interaction effect between Group and Target Hemifield (*F*(2,54)=-.09, *p*=.91).

Taken together, these results show that early target selection processes occur at similar time-points post-stroke, but with reduced magnitude for contralesional stimuli. To examine whether this difference indexes task-relevant behaviour in the two stroke groups, we first calculated asymmetry indices for both RT and N2c amplitude using the formula: (Left Target - Right Target)/((Left Target + Right Target)/2). Note that for N2 asymmetry metrics the denominator was calculated using the absolute value to aid interpretation. Higher indices denote lower amplitudes and slower RTs for left hemifield targets, whereas lower values denote lower amplitudes and slower RTs for right hemifield targets.

Given evidence that the N2c is predictive of RT in younger^19^ and older adults^20^, we explored whether stroke-related changes in this metric were consequential for behaviour. To this end we conducted a multiple regression to examine the relationship between the asymmetry of RT and N2c amplitude in patients, with group included as a covariate. There was a significant positive relationship, whereby higher N2c asymmetries (i.e., lower amplitude for left hemifield targets) were associated with more positive RT asymmetries (i.e., slower RT for left hemifield targets), (F(1,27)=11.84, *p*=.002; Fig. 2; Supplementary Table 2). These data demonstrate that stroke disrupts the strength of early target selection processes in the lesioned hemisphere, and that this effect is relevant to downstream behavioural responses.

### Early Ipsilateral Target Selection Signals Have Lower Ipsilesional Amplitudes After Right Hemisphere Stroke

#### N2i amplitude

To examine group and target hemifield differences in the magnitude of the ipsilateral target selection signal, N2i^19^, we computed an LMM based on single-trial estimates of peak amplitude. There was a significant Group x Target Hemifield interaction (*F*(2,8599)=4.43, *p*=.01; Fig. 2B; Supplementary Table 1). Controls (left: *M*=-6.22 µV/m^2^, *SD*=4.82; right: *M*=-11.43 µV/m^2^, *SD*=4.82) showed a significantly stronger right hemisphere N2i compared with the left hemisphere (*t*(3668)=3.23, *p*corr=.002, model estimate difference=2.38, 95% CI [0.88, 3.88]), consistent with previous work showing that the N2i is right lateralised in healthy individuals^19^. Similar to the controls, the left hemisphere patients had a stronger N2i in the right hemisphere (left: *M*=-5.43 µV/m^2^, *SD*=3.42; right: *M*=-12.47 µV/m^2^, *SD*=3.42); *t*(1800)=4.16, *p* =.001, model estimate difference=3.52, 95% CI [1.53, 5.51]). In contrast to the other two groups, there was no significant difference across hemispheres for the right hemisphere patients (left: *M*=-6.76 µV/m^2^, *SD*=6.67; right: *M*=-6.49 µV/m^2^, *SD*=6.99; *t*(3131)=-0.07, *p*=.94, model estimate difference=-0.078 95% CI [- 2.05, 1.90]). These results thus reflect a lack of right-hemisphere dominance of the N2i that is specific to right hemisphere lesions.

Given this result was exclusive to the right hemisphere patients, we sought to establish whether this was attributable to their marginally (but not significantly) larger stroke-induced lesion volumes, relative to the left hemisphere patients. To this end we performed a multiple regression to examine the relationship between N2i amplitude asymmetry and lesion volume, with group as a covariate. There was no significant relationship between lesion volume and N2i amplitude, *F*(1,18)=0.02, *p*=.89.

Consistent with our approach for the N2c, we next conducted a multiple regression to examine the relationship between the asymmetry of RT and N2i amplitude in our patients, with Group included as a covariate. We did not find evidence for a relationship between the two (*F*(1,27)=0.04, *p*=.85; Supplementary Table 2). Taken together, these results demonstrate that the typical right lateralisation of the N2i is reduced in right hemisphere stroke, but we did not find evidence to suggest that this impacts behaviour.

#### N2i peak latency

We subsequently employed a two-way mixed-effects ANOVA on participant-level means to examine the effects of Group and Target Hemifield on the peak latency of the N2i. There were no significant main effects of Group (*F*(2,54)=0.36, *p*=.70) or Target Hemifield (*F*(1,54)=1.95, *p*=.17), and no significant interaction effect between Group and Target Hemifield (*F*(2,54)=0.14, *p*=.87).

Together, these data suggest that ipsilateral target selection ERPs occur at similar times post-stroke. Right hemisphere strokes are associated with reduced magnitude of these signals in a way that does not appear to meaningfully influence behaviour.

### Evidence Accumulation is Delayed for Stimuli in the Contralesional Hemifield Stimulus-Locked Signal

We extracted the single-trial CPP waveform from central electrode Pz^19,20,42–45^ and defined metrics of time of onset and peak latency (see Online Experimental Procedures). Two-way mixed ANOVAs were then used to investigate differences in onset and peak latency between groups and target hemifields.

#### CPP onset

There was a significant Group x Target Hemifield interaction for CPP onset, *F*(2,53)=11.30, *p*<.001. Bonferroni-Holm adjusted post-hoc analyses revealed that the right hemisphere patients (*t*(16)=3.22, *p*corr<.01) had a later onset of evidence accumulation for left, compared with right, hemifield targets. The left hemisphere patients showed a marginally significant trend in the opposite direction (*t*(11)=-2.45, *p*corr=.06). There were no significant differences between the CPP onsets for left and right hemifield targets for the controls, *t*(26)=-1.31, *p*corr=.20. There were no significant main effects of Group (*F*(2,53)=0.11, *p*=.90) or Target Hemifield (*F*(1,53)=0.60, *p*=.44).

#### CPP peak latency

Similar results were found for the peak latency of the CPP. There were no significant main effects of Group (*F*(2,54)=0.03, *p*=.97) or Target Hemifield (*F*(1,54)=1.26, *p*=.27), but there was a significant interaction (*F*(2,54)=6.34, *p*<.01), again driven by differential target hemifield effects as a function of stroke group. For right hemisphere patients, left hemifield targets elicited a later CPP peak compared with right hemifield targets (*t*(16)=2.69, *p*corr<.05), whereas an opposite trend was apparent in left hemisphere patients (*t*(11)=-2.32, *p*corr=.08). There were no significant differences as a function of target hemifield in the controls (*t*(26)=0.66, *p*corr=.52).

We further examined the relevance of these signals to task-related RT asymmetry exclusively in the stroke groups using two multiple linear regressions (one for each of CPP onset and CPP peak latency asymmetries), with Group included as a covariate. RT asymmetry was related to both CPP onset asymmetry and CPP peak latency asymmetry, such that later onsets (*F*(1,27)=20.76, *p*<.001) and peaks (*F*(1,27)=53.54, *p*<.001) predicted slower behavioural responses.

Overall, evidence accumulation in the stroke groups was slower to begin and end for contralesional, compared with ipsilesional, stimuli. As per our previously published observations in healthy adults^20,43^, markers of delayed evidence accumulation were found to predict slowed responding in both stroke groups (shown Fig. 3).

**Figure 3.**
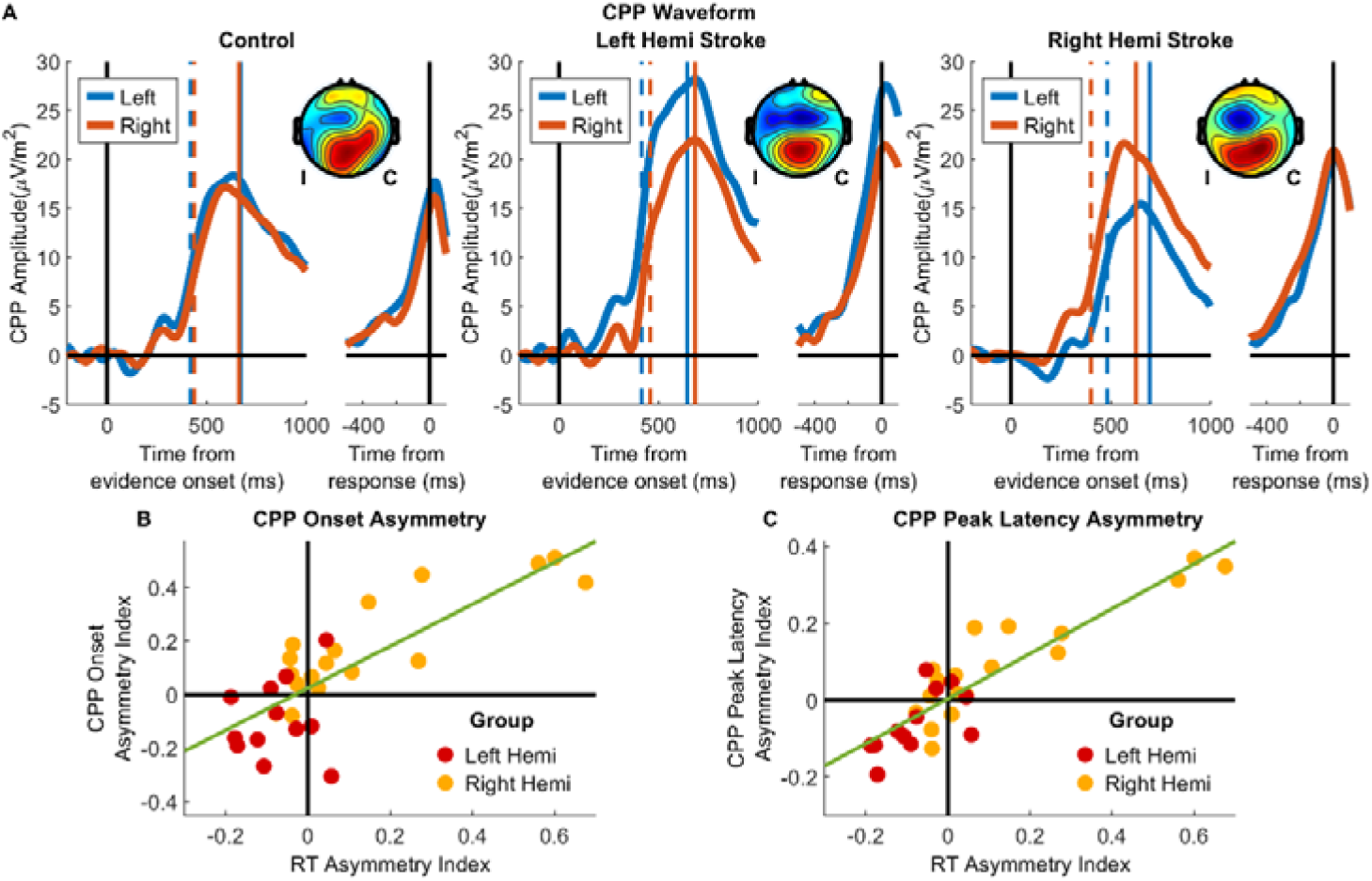
Delayed CPP Onset and Later Peak Latency Index Slowed Contralesional Responding. **A.** Stimulus- and Response-Locked CPP waveforms extracted from electrode Pz, presented separately for each group and for left and right hemifield targets. Inset topoplots reflect average activity between 600-700ms post-stimulus onset for contra- (C) and ipsilateral (I) targets. Dashed lines signify calculated CPP onset, and solid vertical lines indicate CPP peak latency. **B.** Relationship between the CPP onset asymmetry index (where more positive values suggest delayed onset for left hemifield targets) and RT asymmetry index (where more positive values suggest slower responding to left hemifield targets). A least squares regression line is fitted for reference. **C.** Scatter plot of participant means depicting the relationship between CPP peak latency index (where more positive values suggest delayed peaks for left hemifield targets) and RT asymmetry index (where more positive values suggest slower responding to left hemifield stimuli).

#### Response-Locked Signal

To derive a measure of evidence accumulation rate (i.e., CPP slope), we fit a straight line to each participant’s response-locked waveform^18–20,43^ and measured the gradient of the line within a - 150 to 50 ms window around the response^18,20,58^. Peak amplitude was measured as the mean amplitude of the waveform from −50 to 50ms post-response on a per-trial basis^20,42,59^. We then examined group and target hemifield differences in CPP slope using a two-way mixed ANOVA, and peak amplitude using an LMM with fixed effects of Group, Target Hemifield, and a Group x Target Hemifield interaction.

There was no significant main effect of Group (*F*(2,55)=0.17, *p*=.85) or Target Hemifield (*F*(1,55)=1.29, *p*=.26), nor a significant Group x Target Hemifield interaction for the CPP slope (*F*(2,54)=0.73, *p*=.48). Similarly, the interaction effect of Group x Target Hemifield was not significant for CPP peak amplitude (*F*(2,11091)=2.45, *p*=.09), nor were the main effects of Group (*F*(2,11091)=2.42, *p*=.09) or Hemifield (*F*(1,11091)=2.85, *p*=.09).

Overall, stroke did not appear to alter the dynamics of the response-locked CPP, suggesting that evidence is accumulated at similar rates in contralesional and ipsilesional hemispheres.

### Relationship Between Stroke Hemisphere and RT Asymmetry is Mediated by Asymmetries in Target Selection and Evidence Accumulation

The findings above suggest that slowed contralesional responding is indexed by diminished markers of contralateral target selection (N2c) and delayed evidence accumulation (CPP). Given previous work demonstrating that these ERPs represent an interrelated temporal sequence of events^19^, we sought to determine whether they could serially mediate the relationship between stroke hemisphere and RT asymmetry. To test this possibility, we performed a mediation analysis, whereby the degree to which RT asymmetry is predicted by stroke hemisphere is mediated serially by the N2c, followed by the CPP onset. Mediation analyses suggested a full mediation of stroke hemisphere on RT asymmetry via both N2c amplitude and CPP onset asymmetries (indirect effect=0.08, *p*=.04, 95% CI [0.003, 0.15]). A similar mediation was found when the model was repeated with CPP peak latency asymmetry instead of CPP onset (indirect effect=0.14, *p*<.01, 95% CI [.04, .25]). Full mediation results are presented diagrammatically in Fig. 4.

**Figure 4.**
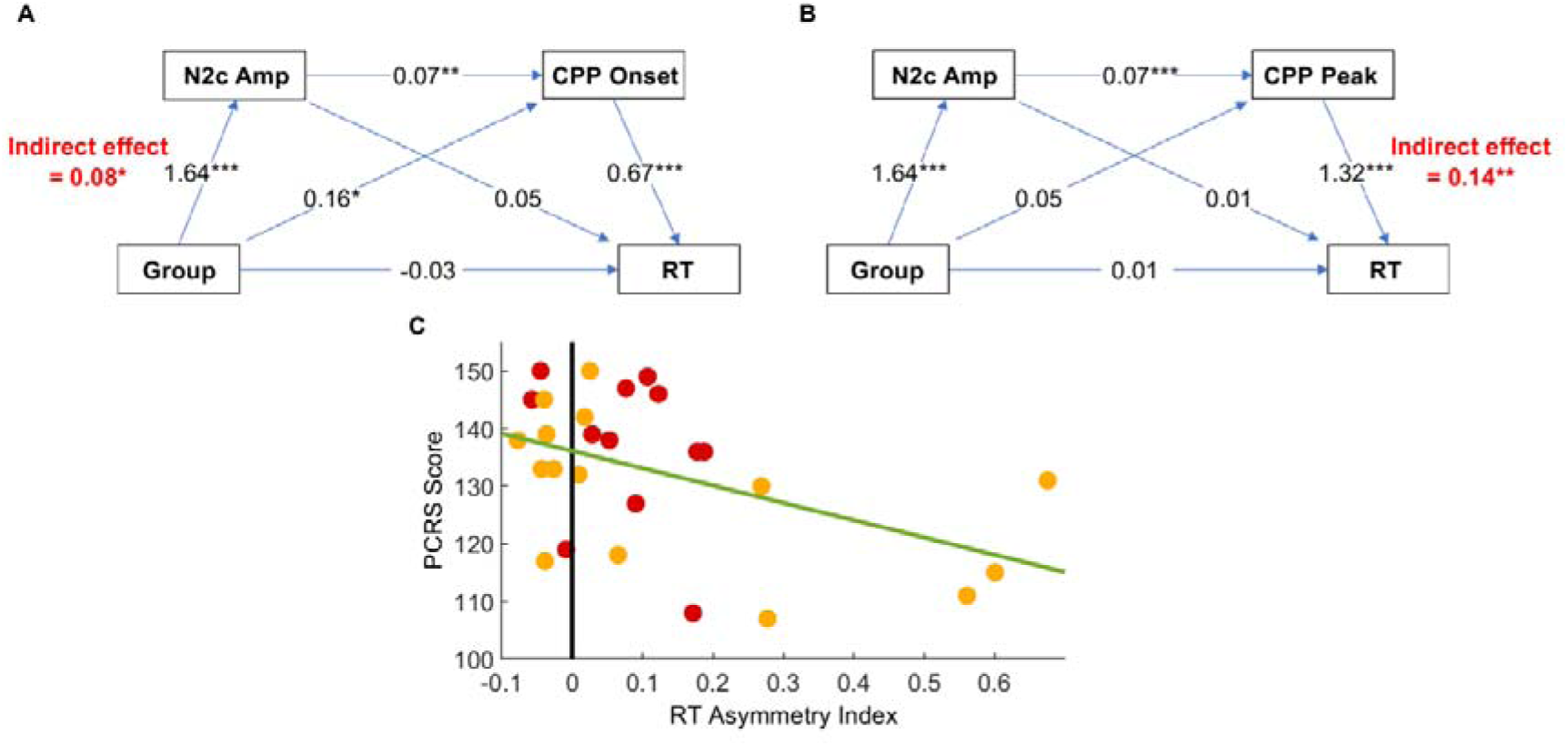
Relationship Between Stroke Side and Contralesional Slowing is Mediated by N2c and CPP Asymmetries. Serial mediation models of lesion side ➔ RT asymmetry including CPP onset latency (**A**) and CPP peak latency (**B**) asymmetries. * *p*<.05, ** *p*<.01, *** *p*<.001. Also depicted is a scatterplot demonstrating the relationship between the self-reported everyday functioning measure (PCRS) and RT asymmetry (**C**).

Collectively, these findings characterise the nature and temporal sequence of discrete neural events that give rise to contralesional slowing after stroke. Specifically, we provide evidence for a mechanism whereby weakened target selection signals (N2c asymmetry) due to stroke delay time-to-threshold of evidence accumulation (CPP onset and peak latency asymmetries) and thence slow contralesional RT.

#### Task-Related Response Speed and Evidence Accumulation Asymmetries Predict Poorer Everyday Functioning

Finally, to determine whether the behavioural asymmetry described above has relevance to daily function following stroke, we asked patients to complete the Patient Competency Rating Scale (PCRS)^60^. The PCRS is a 30-item self-report scale that measures psychosocial functioning post-stroke^61^, whereby greater scores indicate preservation in daily function.^43^

A multiple linear regression examined an asymmetry measure of task RT as a predictor of PCRS scores, with stroke side as a covariate. First, we normalised opposite asymmetries between stroke groups by re-calculating RT asymmetry based on the following formula: (Contralesional Target - Ipsilesional Target)/((Contralesional Target + Ipsilesional Target)/2). This transformation means that greater RT asymmetries indicate slower responses to contralesional stimuli, regardless of lesion side.

Note that there was no significant relationship between lesion volume and RT (*r*=0.17, *p*=.49) or PCRS score (*r*=-.31, *p*=.21) and thus lesion size was not included as a covariate in these analyses.

There was a significant, negative relationship between RT asymmetry on the BRDM task and everyday functioning, such that slower responses to contralesional targets predicted poorer everyday functioning, *F*(1,24)=5.67, *p*=.03. Given this result, we conducted additional, exploratory analyses to examine whether asymmetries in our neurophysiological metrics were related to everyday function. There was marginal evidence for the relevance of CPP parameters to post-stroke daily function, however this was not robust to correction for multiple comparisons (see Supplementary Results 2 and Supplementary Fig. 3).

Taken together, these results suggest that the BRDM task, together with neural metrics of evidence accumulation, are sensitive to lateralised response speed decrements.

## Discussion

Here, we leveraged an integrated EEG/perceptual decision-making paradigm in stroke patients to identify the neurophysiological mechanisms of contralesional slowing in chronic stroke. Results revealed that the established relationship between lesion hemisphere and spatial RT bias was mediated by attenuation of the N2c in response to contralesional stimuli, and subsequently delayed CPP. Importantly, between-hemifield asymmetries in RT were associated with daily functioning. These results therefore suggest similar patterns of neurophysiological change that occur independent of stroke hemisphere and relate to chronic functioning. To our knowledge, this is the first time that a temporal sequence of discrete and well-described neural events such as the target-selective N2c and decision-related CPP has been linked to post-stroke slowing.

These data showed that unilateral lesions selectively weakened the contralateral N2 in the affected hemisphere and subsequently slowed responses. The N2c traces the early selection of goal-relevant targets in uncertain environments and increases proportionally to the quality and availability of sensory evidence^19^, which may suggest that the core deficit following stroke stems from reduced salience of stimuli in the contralesional hemifield. Notably, right hemisphere patients demonstrated additional attenuation of the ipsilateral N2 in the affected hemisphere. Although one might speculate that this may be attributable to the larger average lesion volume in this group such that larger strokes additionally compromised ipsilateral sensory processing, we did not find evidence that this was the case. An alternative explanation is that this effect is specific to lesions in the right hemisphere due to inter-hemispheric differences in the distribution of attention networks (see^62^). There was, however, no evidence that disruptions to ipsilateral target selection meaningfully impacted behaviour, reaffirming the relative importance of contralateral, as opposed to ipsilateral, visual processing^19,44^.

Stroke-related contralesional slowing was additionally correlated with the time of onset and peak of the CPP, an established marker of evidence accumulation^18,20,43,47–56^. Similarly, previous studies on post-stroke contralesional slowing have demonstrated delayed P300^25,31,36^ – an event-related potential that shares many features with the CPP^43,45^. Our findings extend this in three ways. First, our models suggested that the timing of the CPP mediated the relationship between N2c amplitude and RT, and thus demonstrate that delayed evidence accumulation is driven by earlier disruption of target selection, reaffirming the established role of early target selection signals such as the N2c in facilitating and modulating downstream evidence accumulation processes^19^. Second, we show that asymmetry in the timing of evidence accumulation onset directly relates to synchronous performance deficits, increasing the certainty with which these can be hypothesised to be causal. Finally, we demonstrate a dissociation between the time taken to begin evidence accumulation, and the rate at which evidence is sampled: whereas the CPP begins later for contralesional stimuli following stroke, we did not find evidence to suggest that stroke compromises the rate at which the brain accumulates evidence. Thus, following a stroke, decision formation is equally efficient for stimuli presented to ipsilateral and contralesional hemifields, but these processes occur later for contralesional stimuli. Collectively, these data suggest a plausible causal temporal pathway by which unilateral stroke dampens the early selection of sensory stimuli in the contralesional hemifield, and subsequently delays the onset (and thus peak) of evidence accumulation, slowing responses.

Between-hemifield asymmetries in the timing of responses were correlated with daily function, suggesting that the asymmetries displayed here may meaningfully influence behaviour. Asymmetrical sensory processing has been previously shown to deleteriously affect performance of everyday activities such as driving^10^. This may be attributable to the relationship between neural decision-making and higher cognitive functioning^63^ such that hemisphere-specific alterations to the brain’s capacity to select, process and thus respond to relevant stimuli may more broadly impact one’s ability to function within their environment. Exploratory analyses exploring the relationship between the CPP and everyday functioning showed marginal support for this hypothesis, though this effect was not robust to multiple comparisons correction and thus will require replication. Importantly, many of our participants were well beyond the expected critical time points of functional recovery post-stroke^64–67^, indicating that these impacts are persistent and resistant to current rehabilitation methods.

Notably, asymmetries in the neurophysiological signatures of selection and decision processes appeared to be robust determinants of behavioural asymmetries across participants, independent of stroke hemisphere, the presence or absence of overt neglect (though we acknowledge that our use of a single cancellation task is insufficient for comprehensive assessment of neglect, see^68^), and time since stroke. Further, the lesions of our sample were highly heterogeneous with diverse network disconnection profiles and stroke aetiologies, and yet commonalities emerged. Such generalisability speaks to the relevance of neural perceptual decision-making to broader cognition, and the involvement of large, interconnected networks in decision-making^18,63,69^. Collectively, our framework captures behavioural and neural metrics that are relevant to functioning and are generalisable across a wide range of stroke lesions. This is notable as lesion characteristics are often imperfect predictors of outcome, necessitating complex multivariate analyses to achieve some degree of structure-function alignment (see^70^ for discussion of these issues). Given the relevance of these neurophysiological metrics to the behaviour and broader functioning of a wide range of people following stroke, a focus of future work will be to establish whether they have utility in clinical assessment or management over and above that of existing procedures.

In summary, here we apply an integrated EEG paradigm to fractionate the key determinants of pathological response speed biases following stroke. We define a temporal sequence of neurophysiological events from weakened attentional selection in the damaged hemisphere through to delayed evidence accumulation, which together result in slowed perceptual decision-making for contralesional stimuli. This is particularly noteworthy as these prove to be robust neurophysiological predictors of behaviour despite the heterogeneity of our sample’s lesion characteristics. Taken together, our results underscore the potential utility of neurophysiological assessment for describing clinically relevant behaviour.

## Experimental Procedures (Online Only)

### Participants

The experimental protocol was approved by the Oxford Central University (MSD IDREC reference R61968/RE001), Monash Health (MHREC reference #14408A) and Monash University Human Research Ethics Committees (MUHREC reference #2492) prior to the commencement of the study. All participants volunteered without knowledge of the hypotheses being tested and provided written informed consent prior to participation.

### Neurologically Healthy Controls

A total of 31 neurologically healthy adults aged between 57 and 90 years were recruited from the community in Melbourne, Victoria, Australia. All participants reported normal or corrected-to-normal visual acuity, self-reported being right-handed in all common daily tasks (*n*=1 excluded due to left handedness), were not taking arousal-modulating medications (such as tranquilisers or steroids) and reported no significant previous neurological or psychiatric history (*n*=1 excluded due to extensive medical history). All participants were required to score higher than 23/30 on the Montreal Cognitive Assessment (MoCA)^71^ after adjusting for education ^97^ (*n*=1 excluded). One further participant was excluded due to EEG data quality issues, leaving a total of *n*=27 controls for analysis.

### Post-Stroke Adults

Seventy-one (71) individuals who had experienced a focal, first-ever clinically significant unilateral cerebral stroke were recruited for the study. Focusing exclusively on unilateral stroke enabled us to consider the specific role of each hemisphere in regulating behaviour. Of these, 38 were recruited from Melbourne, Victoria, Australia (*n*=8 from an existing Monash University registry; *n*=22 directly from Monash Health subacute inpatient rehabilitation facilities at the Kingston Centre, Cheltenham; and *n*=8 via letters sent to patients discharged from Monash Health facilities between 08/09/2019 and 08/09/2021). A further 33 patients were recruited from an existing stroke research volunteer database managed by the Translational Neuropsychology Group, University of Oxford, United Kingdom. All participants were right-handed prior to their stroke, had a clinically confirmed medical diagnosis of stroke, had no prior history of significant neurological or psychiatric disorder other than stroke, reported normal or corrected-to-normal visual acuity, and had no reported use of arousal-altering medications. Participants did not have visual field defects that could otherwise compromise task performance (i.e., not in the inferior quadrants where the visual stimuli were presented), as measured by a simple visual field assessment (within the Oxford Cognitive Screen; OCS^72^), and did not have a primary extinction syndrome on a computerised extinction task^73^.

Of these, *n*=55 patients completed the study (*n*=1 was deemed ambidextrous pre-stroke; *n*=12 were unable to complete the task due to cognitive difficulties (i.e., marked disinhibition leading to excessive false alarms or comprehension difficulties) or difficulties maintaining central fixation; *n*=3 were lost to attrition. A further *n*=21 were later excluded following medical record and brain imaging review (*n*=1 had their diagnosis updated to a transient ischaemic attack; *n*=11 had neuroimaging suggestive of bilateral lesions; *n*=4 had multi-focal strokes; *n*=2 had records suggestive of post-stroke epilepsy; *n*=2 had substantial unreported medical histories; *n*=1 had cognitive complaints that pre-dated their stroke). Finally, *n*=4 were removed due to data quality issues such as low trial counts or extremely noisy EEG data, leaving a total of *n*=30 participants. We note that of these, *n*=2 participants used a left-hand response due to right-sided hemiplegia, and *n*=3 had upper quadrant visual field defects. These participants were not excluded as the visual stimuli in the random-dot motion task were presented in the lower half of the screen. Enforced central fixation using real-time pupil recordings ensured that the bilateral visual stimuli were always displayed exclusively within the left and right visual hemifields.

Demographic information for the entire sample of *N*=57 participants is presented in Table 1. Clinical stroke characteristics are displayed in Table 2.

**Table 1.**
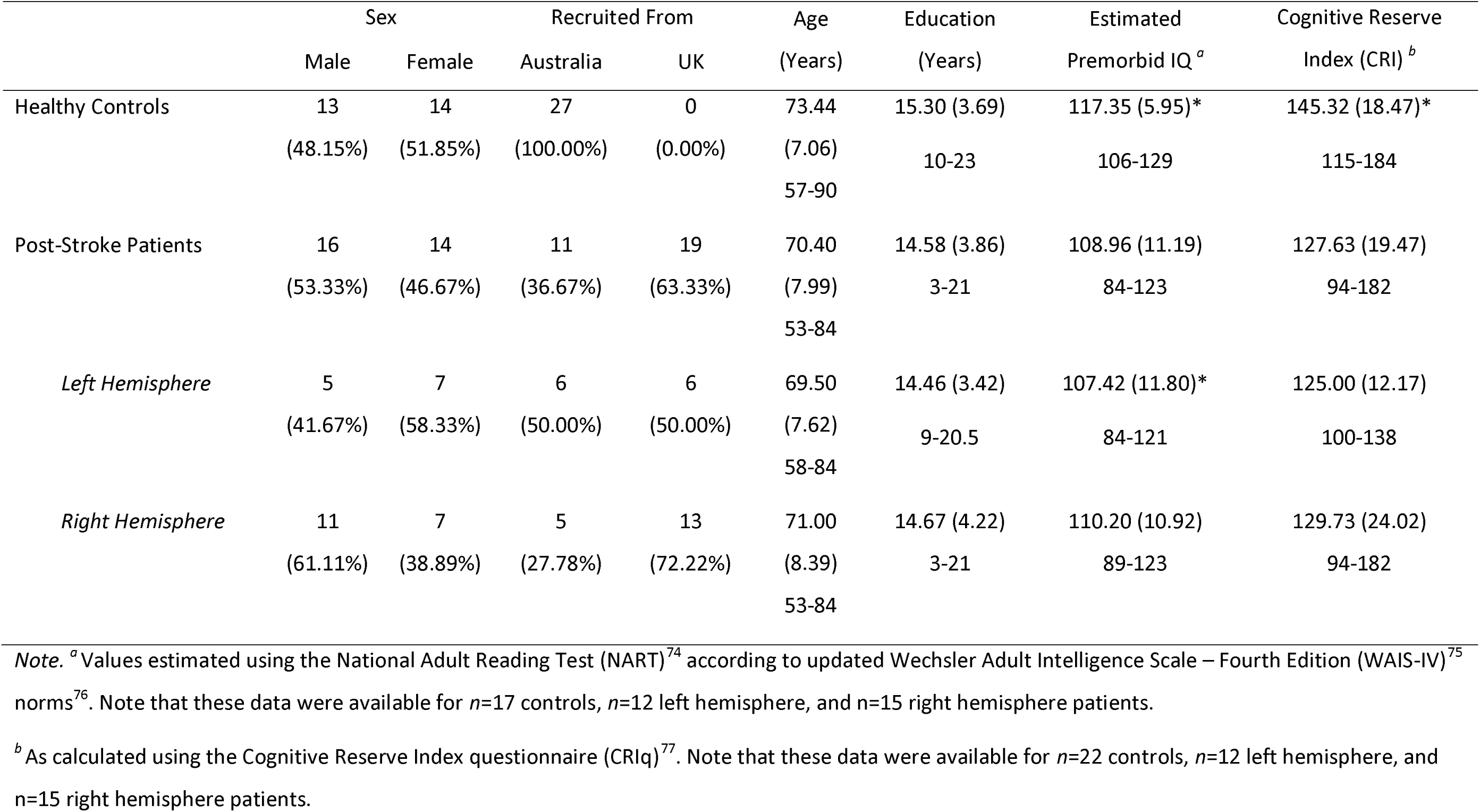
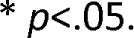
Demographic Information Across.

|  | Sex |  | Recruited From |  | Age | Education | Estimated | Cognitive Reserve |
| --- | --- | --- | --- | --- | --- | --- | --- | --- |
|  | Male | Female | Australia | UK | (Years) | (Years) | Premorbid IQ <sup>a</sup> | Index (CRI) <sup>b</sup> |
| Healthy Controls | 13<br>(48.15%) | 14<br>(51.85%) | 27<br>(100.00%) | 0<br>(0.00%) | 73.44<br>(7.06)<br>57-90 | 15.30 (3.69)<br>10-23 | 117.35 (5.95)*<br>106-129 | 145.32 (18.47)*<br>115-184 |
| Post-Stroke Patients | 16<br>(53.33%) | 14<br>(46.67%) | 11<br>(36.67%) | 19<br>(63.33%) | 70.40<br>(7.99)<br>53-84 | 14.58 (3.86)<br>3-21 | 108.96 (11.19)<br>84-123 | 127.63 (19.47)<br>94-182 |
| *Left Hemisphere* | 5<br>(41.67%) | 7<br>(58.33%) | 6<br>(50.00%) | 6<br>(50.00%) | 69.50<br>(7.62)<br>58-84 | 14.46 (3.42)<br>9-20.5 | 107.42 (11.80)*<br>84-121 | 125.00 (12.17)<br>100-138 |
| *Right Hemisphere* | 11<br>(61.11%) | 7<br>(38.89%) | 5<br>(27.78%) | 13<br>(72.22%) | 71.00<br>(8.39)<br>53-84 | 14.67 (4.22)<br>3-21 | 110.20 (10.92)<br>89-123 | 129.73 (24.02)<br>94-182 |

*Note.* <sup>a</sup> Values estimated using the National Adult Reading Test (NART)<sup>74</sup> according to updated Wechsler Adult Intelligence Scale – Fourth Edition (WAIS-IV)<sup>75</sup> norms<sup>76</sup>. Note that these data were available for $n=17$ controls, $n=12$ left hemisphere, and $n=15$ right hemisphere patients.
<sup>b</sup> As calculated using the Cognitive Reserve Index questionnaire (CRIq)<sup>77</sup>. Note that these data were available for $n=22$ controls, $n=12$ left hemisphere, and $n=15$ right hemisphere patients.
\* $p < .05$ .

**Table 2.**
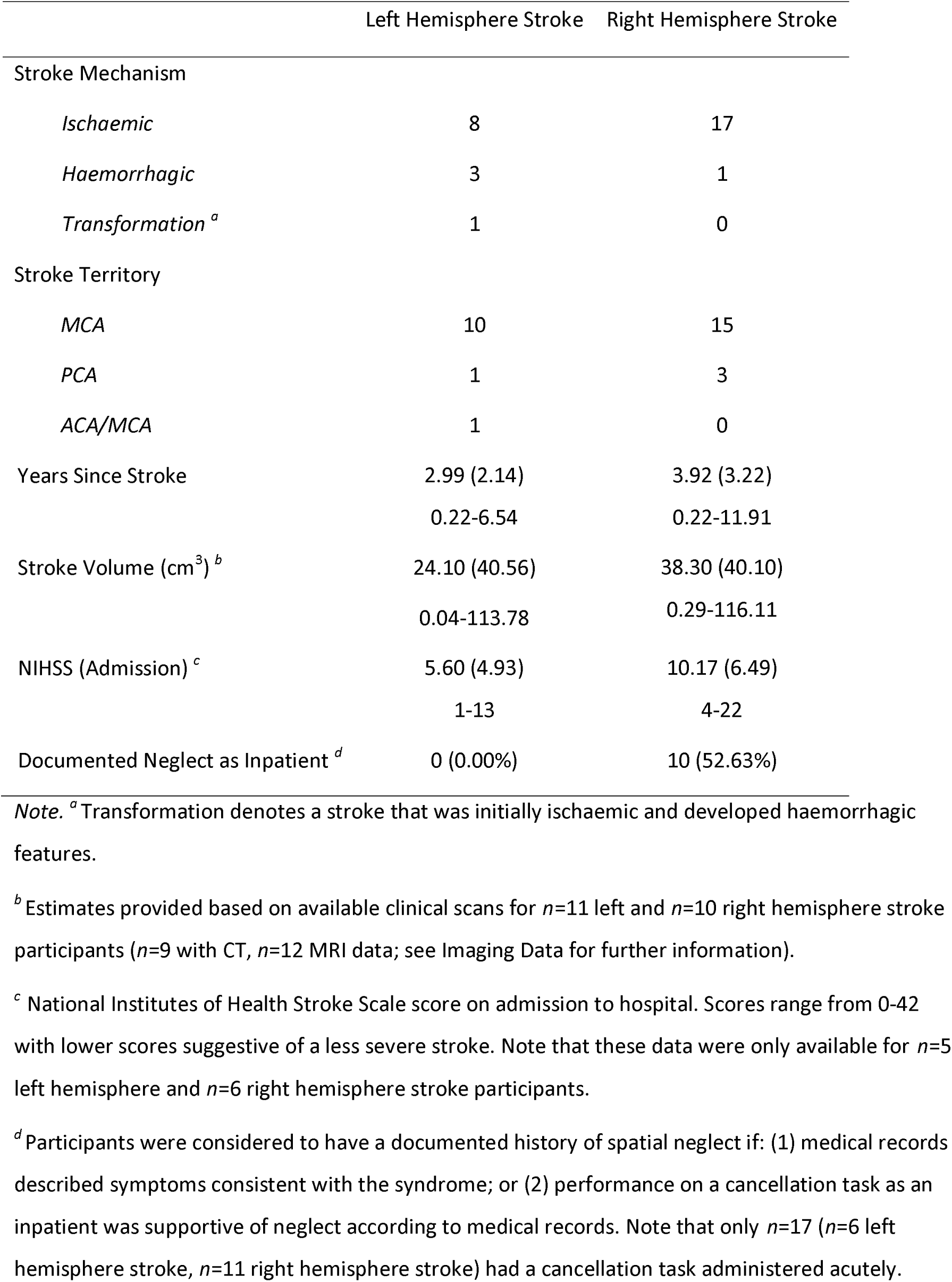
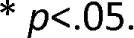
Stroke Characteristics.

|  | Left Hemisphere Stroke | Right Hemisphere Stroke |
| --- | --- | --- |
| Stroke Mechanism |  |  |
| *Ischaemic* | 8 | 17 |
| *Haemorrhagic* | 3 | 1 |
| *Transformation*<sup>a</sup> | 1 | 0 |
| Stroke Territory |  |  |
| *MCA* | 10 | 15 |
| *PCA* | 1 | 3 |
| *ACA/MCA* | 1 | 0 |
| Years Since Stroke | 2.99 (2.14)<br>0.22-6.54 | 3.92 (3.22)<br>0.22-11.91 |
| Stroke Volume (cm<sup>3</sup>)<sup>b</sup> | 24.10 (40.56)<br>0.04-113.78 | 38.30 (40.10)<br>0.29-116.11 |
| NIHSS (Admission)<sup>c</sup> | 5.60 (4.93)<br>1-13 | 10.17 (6.49)<br>4-22 |
| Documented Neglect as Inpatient<sup>d</sup> | 0 (0.00%) | 10 (52.63%) |

*Note.* <sup>a</sup> Transformation denotes a stroke that was initially ischaemic and developed haemorrhagic features.
<sup>b</sup> Estimates provided based on available clinical scans for $n=11$ left and $n=10$ right hemisphere stroke participants ( $n=9$ with CT, $n=12$ MRI data; see Imaging Data for further information).
<sup>c</sup> National Institutes of Health Stroke Scale score on admission to hospital. Scores range from 0-42 with lower scores suggestive of a less severe stroke. Note that these data were only available for $n=5$ left hemisphere and $n=6$ right hemisphere stroke participants.
<sup>d</sup> Participants were considered to have a documented history of spatial neglect if: (1) medical records described symptoms consistent with the syndrome; or (2) performance on a cancellation task as an inpatient was supportive of neglect according to medical records. Note that only $n=17$ ( $n=6$ left hemisphere stroke, $n=11$ right hemisphere stroke) had a cancellation task administered acutely.
\* $p < .05$ .

### Procedure & Materials

All participants completed between one and four testing sessions (mean=2.87, median=3.00), depending on fatigue, during which they completed the tasks described below, plus an additional computerised measure not considered here. Testing took place at either Monash University, Clayton, Victoria, Australia, or Oxford University, Oxford, England. Both controls and patients completed the Bilateral Random Dot Motion (BRDM) task^19,20^ (Fig. 1) while being monitored with EEG. Eye movements were monitored with an EyeLink 1000 remote infrared eye-tracker (SR Research Ltd). The task was controlled via a Windows PC using MATLAB (MathWorks) and the Psychophysics Toolbox extensions^78–80^, and stimuli were displayed on either a 21-inch CRT monitor (Melbourne), or on a 23- or 27-inch LED monitor (Oxford). All monitors were set to a refresh rate of 85Hz and a screen resolution of 1024 x 768. Participants viewed the task from 57cm in a darkened, sound-attenuated room, with their head supported by a chin rest.

Prior to completion of the BRDM EEG task, stroke patients completed a larger battery of cognitive tasks including the OCS^72^, the Auditory Attention Task (AAT) from the Birmingham Cognitive Screen (BCoS)^81^, a Landmark task (e.g.^82^), and a self-report questionnaire, the Patient Competency Rating Scale (PCRS)^60^. This questionnaire was completed with the assistance of close others where insight was compromised. Note that to minimise interference with ongoing treatment and rehabilitation, the subset of n=2 included stroke participants recruited from acute stroke wards did not complete the cognitive tasks or questionnaires.

### Behavioural Assessment of Perceptual Decision-Making

The bilateral random dot motion task^19,83^ is a computerised perceptual decision-making task that allows fine-grained examination of how quickly and accurately responses are made separately for targets in each visual hemifield. Participants were required to maintain central fixation while monitoring two peripheral circular patches that were positioned 10° either side and 4° below the fixation square, covering 8° visual angle (see Fig. 1). These patches consisted of 150 9×9 pixel randomly moving dots which were displaced by five degrees per second. At intervals of 1.8s, 2.8s, or 3.8s, 90% of the dots in either patch transitioned to moving uniformly either upward or downward. These target events occurred with equal probability in the left and right patches. Participants were instructed to respond via a speeded single mouse click with their right index finger (left index finger for those with right-sided hemiplegia, n=2) as soon as they detected any coherent motion targets. Note that trials restarted if a blink or gaze deviation more than four degrees left or right of centre was detected, as monitored by an EyeLink 1000 remote infrared eye-tracker (SR Research Ltd). Coherent motion pulses persisted for 3s or until a response was made.

Practice trials were administered until the participant understood the task, irrespective of their accuracy on these trials. Participants then completed 7 – 15 blocks of 24 trials. There were 12 possible trial types which varied according to the inter-stimulus interval (1.8s, 2.8s, or 3.8s), hemifield of target (left or right), and motion direction (up or down), with each presented twice per block in pseudorandom order. Each block was followed by breaks of up to 180 seconds.

### Cognitive Measures

#### Oxford Cognitive Screen (OCS)

The OCS^72^ is a stroke-specific, aphasia- and neglect-inclusive cognitive screening tool initially designed for use in acute settings. It features ten separate tasks that capture the domains of language, praxis, numeracy, memory, visuospatial function, and attention. In particular it features an embedded cancellation task (‘Broken Hearts’) which has an overall sensitivity of 94.12% for spatial neglect, as compared with the star cancellation task^72^. The OCS has been demonstrated to have superior sensitivity in detecting post-stroke cognitive impairment relative to other common screening tasks, and has been recommended for first-line neglect screening^84,85^.

#### Auditory Attention Task (AAT)

The AAT is a subtest of the BCoS^81^, a screening task designed for use in acquired brain injury. The AAT measures sustained attention by asking participants to listen to a stream of words and tap on the table whenever they hear one of three target words. This allows a measure of both omissions (i.e., missed responses) and false positives (i.e., impulsive responses to non-target words). The BCoS itself demonstrates good psychometric properties^81^ and has utility in eliciting post-stroke changes in cognition^86^.

#### Landmark task

The Landmark task (e.g.,^82^) was included as a measure of visuospatial bias. Participants were supplied with 20 bisected horizontal lines and asked to indicate whether the left or right side of the line was shorter. In ten trials the line is bisected centrally, and in the remainder the bisection is either offset to the left (five trials) or right side (five trials). On each side, three trials are offset by 1mm, by 2.5mm, and by 5mm. We calculated spatial bias using only the ten centrally-bisected trials with the formula: (left responses – right responses)/10. Possible scores therefore ranged from −1 to 1, with more negative scores suggestive of left spatial bias (i.e., the participant reported more of the centrally bisected lines were shorter on the right side) and more positive scores indicative of right spatial bias. Scores of zero suggest no consistent bias.

### Functional Questionnaires

#### Patient Competency Rating Scale (PCRS)

Self-reported psychosocial functioning was examined using the 30-item PCRS^60^. Participants rated the ease with which they could perform a range of common activities (e.g., dressing themselves, starting a conversation in a group, or remembering their daily schedule) on a five-point Likert scale from 1 (“Can’t do”) to 5 (“Can do with ease”). Scores were aggregated with possible scores ranging from 30 to 150. Higher scores are suggestive of higher functioning. Although initially designed to measure insight following traumatic brain injury, the scale has been validated for characterising psychosocial functioning following stroke^61^.

Descriptive data for the cognitive tasks and questionnaires are presented in Table 3.

**Table 3.**
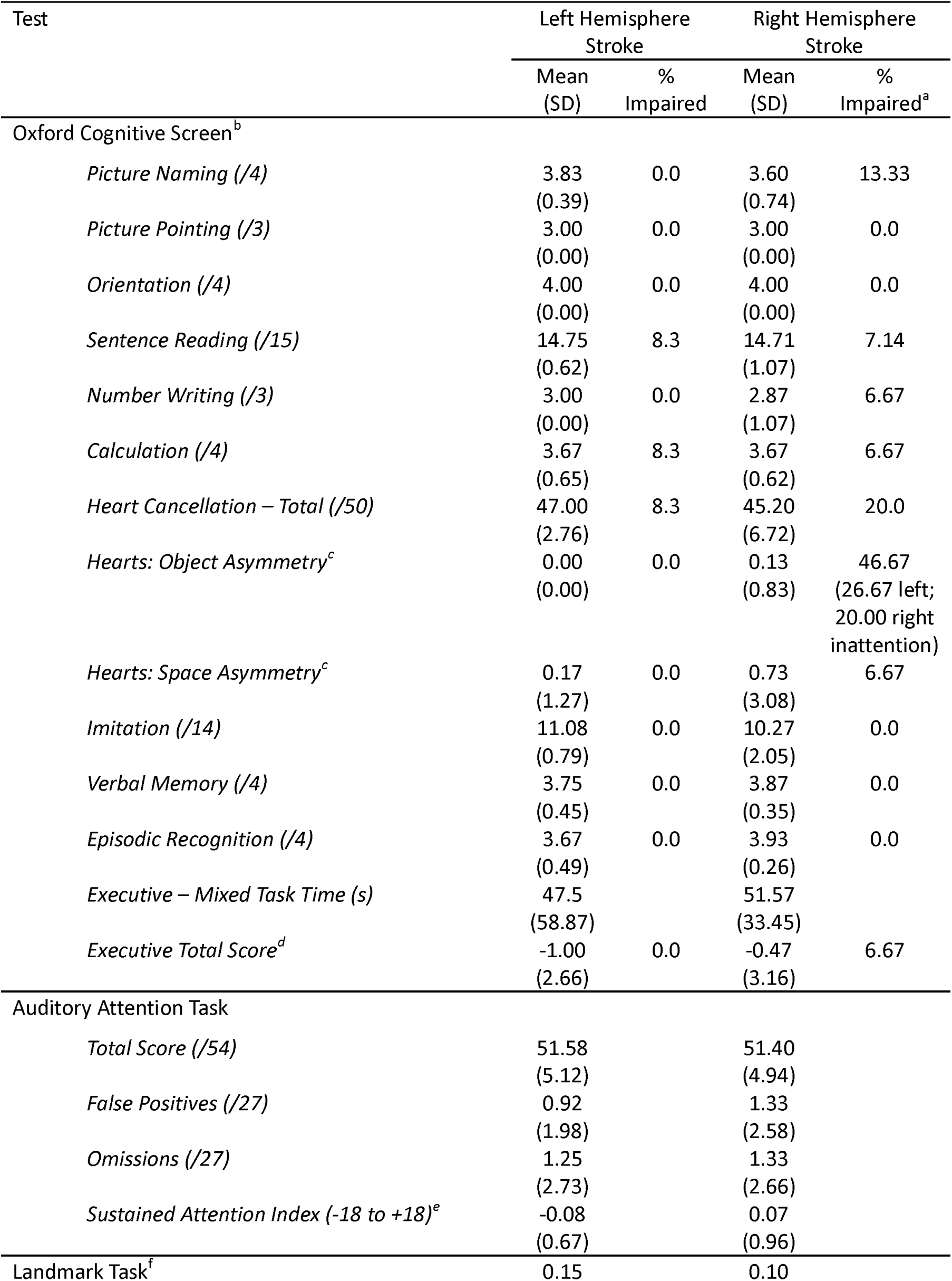
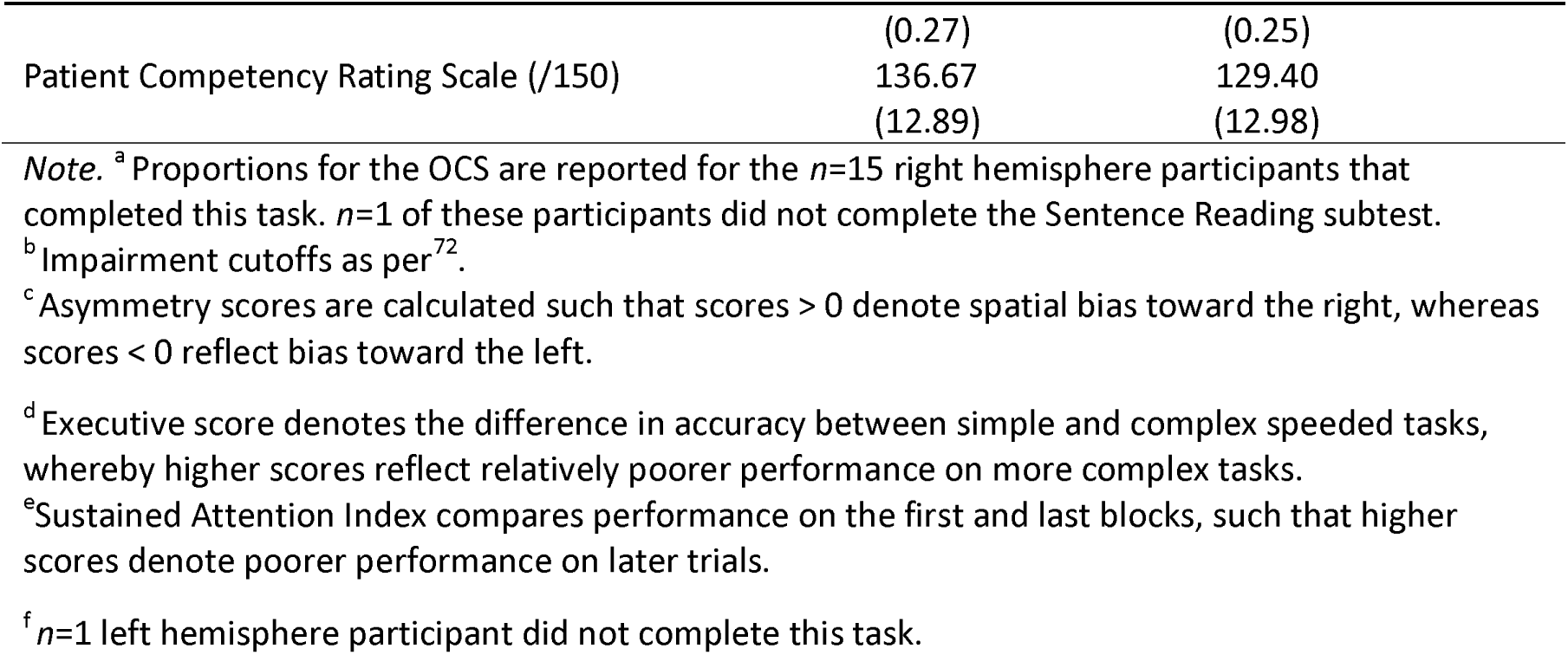
Cognitive Test Data by Group.

#### Imaging Data

Where possible we acquired clinical CT (n=9) and/or MRI scans (n=12) for the final subset of participants. For those with MRI brain data, we manually selected the image that best demonstrated the lesion (1 DWI, 7 FLAIR, 2 T2, 2 T1). In the remaining n=9 participants, no imaging was available. Recent research has verified that similar routine clinical imaging data are of sufficient quality to accurately localise established neural correlates in statistical lesion mapping analyses^87,88^. Prior to any behavioural analysis, lesion masks were derived and processed in line with a standard analysis pipeline^89^. Specifically, lesions were manually delineated on the axial plane of native space scans by a trained expert (MJM) using MRIcron^90^. These native-space masks were smoothed at 5mm full width at half maximum in the z-direction, reoriented to the anterior commissure and normalised to 1×1×1 MNI space. This spatial normalisation was completed using Statistical Parametric Mapping^91^ and standard, age-matched templates (Clinical Toolbox)^92^. All normalised scans and lesions were visually inspected to ensure accuracy prior to analysis.

As the lesion data available for the study do not meet the recommended lesion overlay statistical power minimum for statistical lesion mapping analyses^93^, quantitative lesion mapping analyses were not conducted. However, lesion overlay comparisons were conducted to evaluate broad, qualitative relationships between lesion location and behavioural/EEG metrics. For each relevant metric, lesion maps were colour coded to represent behavioural/EEG scores. All masks were then overlayed in MNI space to visualise lesion-behaviour relationships. Lesion maps are presented for each stroke group in Fig. 1.

To quantify network-level disconnection, lesion masks were down sampled onto the Schaefer-Yeo Atlas Parcellation (100 parcels, 7 Networks)^94^. To estimate parcel-wise disconnection severity, the percentage of disconnected HCP-842 streamlines that bilaterally terminate within each pair of included grey matter parcels was calculated using the Lesion Quantification Toolkit^95^. Additional subcortical and cerebellar parcels derived from the AAL^96^ and Harvard-Oxford Subcortical Atlas, respectively, were also included (n=35). Individual lesion masks, network disconnection summaries, and cortical integrity statistics can be found in Extended Data 1, 2, and 3, respectively.

#### Electroencephalography (EEG)

Continuous EEG were acquired from either 64 scalp electrodes using a BrainAmp DC system (Brain Products), digitised at 500Hz (Melbourne), or 64 electrodes using a Neuroscan SynAmpsRT system, digitised at 1000Hz (Oxford), while participants completed the BRDM. We used a combination of custom scripts (available at https://github.com/gerontium/AusOxStroke) and EEGLAB routines^97^ implemented in MATLAB (MathWorks) to process the data.

#### Pre-Processing

First, we manually inspected channel variances to identify channels artifacts for later interpolation. We then accounted for cross-site differences in data collection by re-aligning the channels according to a common subset of 58 electrodes. Data collected in Oxford was re-sampled from 1000 to 500 Hz. Thereafter all data were treated consistently. Data were detrended and we eliminated line noise by applying notch filters at 50, 100, and 150 Hz. We then applied a 0.1 Hz Hamming windowed sinc FIR high-pass filter. Channels previously identified as demonstrating zero or extreme variance were interpolated via spherical spline (mean channels interpolated = 1.95, median = 0). The interpolated data were then low-pass filtered at 8 Hz (120 Hz for spectral analyses) via similar methods, and re-referenced to the average reference.

A trial was defined as the period between central fixation and either a response, a fixation break, or onset of new random motion stimulus, as identified using EEG-recorded triggers. We extracted epochs from −200ms pre-target onset to 3000ms post-target onset (i.e., target offset), while response-locked epochs were extracted from −500ms pre-response to 100ms post-response. We then removed trials meeting any of the following conditions: (1) one or more EEG channels exceeded 100 µV between baseline (200 ms pre-target onset) and 100ms post-response, or between baseline and 1000 ms post-target onset; or (2) the participant blinked or deviated their gaze by more than 3° left or right of centre during the same time window. Finally, we applied a Current Source Density transformation with a spline flexibility of 4 to disentangle overlapping ERP components^98^, and baselined the stimulus- and response-locked ERP waveforms relative to −200 to 0 ms pre-stimulus onset.

#### Event-Related Potential (ERP) Extraction

CPP and N2 waveforms were derived by aggregating the baseline-corrected epochs at relevant electrodes separately for left and right hemifield targets. Consistent with our previous publications^18–20,44^, we extracted the N2c contralateral to target location at peak electrodes P8 and P7 for left and right hemifield targets, respectively. The ipsilateral N2i component was extracted at peak electrodes P8/TP8 and P7/TP7. The CPP was measured centrally at peak electrode Pz irrespective of target hemifield^19,20,42–45^.

We subsequently derived a number of metrics for each of the stimulus-locked ERP waveforms on a single-trial basis for each participant. The peak latency of the N2c was derived according to the mean stimulus-locked waveform for each participant, and was defined as the time point that reflected the most negative amplitude value between 150-400 ms post-stimulus onset^19,20^. We calculated N2c amplitude as the mean amplitude within a 10 ms window centred on the N2c’s grand average peak^19,20^. The same was done to calculate peak amplitude and latency of the N2i, but from ipsilateral electrodes. To increase the accuracy of N2c/N2i amplitude and decrease any influence from slow drifts, we subtracted a baseline amplitude up to 180 ms post-target, given the lack of ERP response in that period^19^. To derive measures of CPP onset and peak latency, we first applied a 2 Hz low-pass filter to the stimulus-locked CPP waveform to smooth out noise that might interfere with these calculations. Peak latency was measured as the time point at which the CPP waveform from 400-1000 ms (the end of the EEG artifact rejection window) post-stimulus onset reached its peak amplitude. CPP onset was defined as the timepoint at which the CPP waveform reached half of its peak amplitude.

Additional neural metrics of build-up rate and peak amplitude were calculated using the response-locked CPP waveform. We first fitted a straight line to each participant’s waveform^18–20,43^ and defined the CPP build-up rate as the slope within a −150 to −50 ms window around the response^18,20,58^. Finally, CPP amplitude was defined as the mean amplitude of the response-locked waveform from −25 to 25ms post-response^20,42,59^.

### Statistical Analyses

Many of the conducted analyses utilised the same structure, investigating the existence of a Group (Healthy, Left Stroke, Right Stroke) x Target Hemifield (Left, Right) interaction. For RT, N2c amplitude, N2i amplitude and CPP peak amplitude, we took advantage of single trial analysis using Linear Mixed Effects (LME) modelling, predicting the given outcome measure with fixed effects of Group and Target Hemifield and random slopes and intercepts for Target Hemifield x Participant. Thus, the equation for those models was the following: [RT/N2c/N2i/CPP] ∼ TargetHemifield x Group + (TargetHemifield|Participant). This was the maximal model that could be fitted and permitted individual participant effects to vary, while still revealing group-level effects. Other analyses were conducted on a participant-level to increase signal to noise ratio before measurement; N2c peak latency, N2i peak latency, CPP onset latency, CPP peak latency and CPP build-up rate. These analyses were performed using a mixed-effects ANOVA, with factors of Target Hemifield and Group. When using LME, we performed Bonferroni-Holm-adjusted post-hoc analyses comparing Target Hemifield within each Group using further LMEs, whereas when using ANOVA, we performed post-hoc analyses using within group t-tests and calculated Bonferroni-Holm-adjusted p values to account for multiple comparisons.

Next, for Left and Right Stroke Groups, we calculated Left Hemifield minus Right Hemifield asymmetry indices for both RT and our ERP metrics of interest via the following formula: (Left Target - Right Target)/((Left Target + Right Target)/2). Thus, for RT and ERP latency variables, lower indices denoted earlier timing to left hemifield targets, whereas higher indices reflected earlier timing for right hemifield targets. There was one exception to this strategy for N2c/N2i amplitude asymmetry indices, which were calculated using the following formula: (Left Target - Right Target)((abs(Left Target) + (abs(Right Target))/2). Note the absolute of the signals was used in the denominator for these indices, due the fact that the N2 signals could be negative. For these amplitude metrics, lower indices were indicative of greater (i.e., more negative) N2c/N2i amplitude in response to left hemifield targets, whereas higher indices denoted greater amplitude in response to right hemifield targets. To ascertain the relationship between these indices, we performed a linear regression, covarying for Stroke Group to ensure that effects were not simply driven by differences between groups.

We followed up these analyses with two serial mediation analyses to investigate whether the effect of Stroke Group on RT asymmetry was mediated via N2c amplitude asymmetry, then CPP onset (and peak latency in a secondary mediation) asymmetry. To model this, we performed two, two-stage serial mediations, with Stroke Group as predictor, then N2c amplitude asymmetry and CPP onset/CPP peak latency asymmetry as serial mediators, and RT asymmetry as the Dependent Variable.

Finally, we tested the relationship between RT asymmetry and Quality of Life indices, measured by the PCRS. Given that PCRS is an absolute measure, we calculated a slightly different RT asymmetry index which denoted worse contralesional RTs: (Contra Target - Ipsi Target)((Contra Target + Ipsi Target)/2). A lower index here denoted faster RT to targets contralateral to the lesioned hemisphere, whereas a higher index denoted slower RT to targets contralateral to the lesioned hemisphere, i.e., the effect one might expect post-stroke. We then performed a linear regression of RT asymmetry index on PCRS, covarying for Stroke Group.

## Acknowledgements

This work was funded by an NHMRC Project Grant to MAB, JBM and RGO (#1140707); Australian Research Council Grants DP150100986 and DP180102066 to M.A.B and R.G.O; a European Commission Grant #844246 and a Cullen Junior Research Fellowship at Corpus Christi College, University of Oxford, to M.B.B.; a National Institute for Health Research Oxford Health Biomedical Research Center and Wellcome Centre for Integrative Neuroimaging Wellcome Trust Grant 203139/Z/16/Z; an Australian NHMRC Senior Research Fellowship (Level B; APP1154378) to M.A.B.; a European Research Council Grant #865474 to R.G.O.; an Australian Research Council Grant FTFT220100294 to T.T.-J.C.; a National Institute for Health Research Advanced Fellowship NIHR302224 to N.D.; an NHMRC Investigator Grant (Leadership 3; GNT2010141) to J.B.M.; an NHMRC Ideas grant (APP1186955) to D.R.; and an Australian Government Research Training Program (RTP) Scholarship to D.J.P.

## References

1. Feigin, V. L., et al. Global, regional, and national burden of stroke and its risk factors, 1990-2019: a systematic analysis for the Global Burden of Disease Study 2019. Lancet Neurol 20, 1–26 (2021).

2. Ballard, C., et al. Profile of Neuropsychological Deficits in Older Stroke Survivors without Dementia. Dement Geriatr Cogn Disord 16, 52–56 (2003).

3. Hurford, R., Charidimou, A., Fox, Z., Cipolotti, L. & Werring, D. J. Domain-specific trends in cognitive impairment after acute ischaemic stroke. J Neurol 260, 237–241 (2013).

4. Johansson, B. & Rönnbäck, L. Mental fatigue and cognitive impairment after an almost neurological recovered stroke. ISRN Psychiatry 2012, 1–7 (2012).

5. Losier, B. J. W. & Klein, R. M. A review of the evidence for a disengage deficit following parietal lobe damage. Neurosci Biobehav Rev 25, 1–13 (2001).

6. Pihlaja, R., Uimonen, J., Mustanoja, S., Tatlisumak, T. & Poutiainen, E. Post-stroke fatigue is associated with impaired processing speed and memory functions in first-ever stroke patients. J Psychosom Res 77, 380–384 (2014).

7. Su, C. Y., Wuang, Y. P., Lin, Y. H. & Su, J. H. The Role of Processing Speed in Post-Stroke Cognitive Dysfunction. Archives of Clinical Neuropsychology 30, 148–160 (2015).

8. Mahon, S. et al. Slowed Information Processing Speed at Four Years Poststroke: Evidence and Predictors from a Population-Based Follow-up Study. Journal of Stroke and Cerebrovascular Diseases 29, 104513 (2020).

9. Gerritsen, M. J. J., Berg, I. J., Deelman, B. G., Visser-Keizer, A. C. & Meyboom-De Jong, B. Speed of Information Processing After Unilateral Stroke. J Clin Exp Neuropsychol 25, 1–13 (2003).

10. Van Kessel, M. E., Van Nes, I. J. W., Geurts, A. C. H., Brouwer, W. H. & Fasotti, L. Visuospatial asymmetry in dual-task performance after subacute stroke. J Neuropsychol 7, 72–90 (2013).

11. Stephens, S. et al. Association between mild vascular cognitive impairment and impaired activities of daily living in older stroke survivors without dementia. J Am Geriatr Soc 53, 103– 107 (2005).

12. Salthouse, T. A. The processing-speed theory of adult age differences in cognition. Psychol Rev 103, 403–428 (1996).

13. Kail, R. & Salthouse, T. A. Processing speed as a mental capacity. Acta Psychol (Amst) 86, 199– 225 (1994).

14. Sterzer, P. Moving forward in perceptual decision making. Proc Natl Acad Sci U S A 113, 5771– 5773 (2016).

15. Imani, E., Harati, A., Pourreza, H. & Goudarzi, M. M. Brain-behavior relationships in the perceptual decision-making process through cognitive processing stages. Neuropsychologia 155, 107821 (2021).

16. Hauser, C. K. & Salinas, E. Perceptual Decision Making. Encyclopedia of Computational Neuroscience 1–21 (2014) doi:10.1007/978-1-4614-7320-6_317-1.

17. Brooks, J. X. & Cullen, K. E. Predictive Sensing: The Role of Motor Signals in Sensory Processing. Biol Psychiatry Cogn Neurosci Neuroimaging 4, 842–850 (2019).

18. Brosnan, M. B. et al. Evidence accumulation during perceptual decisions in humans varies as a function of dorsal frontoparietal organization. Nature Human Behaviour 2020 4:8 4, 844–855 (2020).

19. Loughnane, G. M. et al. Target Selection Signals Influence Perceptual Decisions by Modulating the Onset and Rate of Evidence Accumulation. Curr Biol 26, 496–502 (2016).

20. Brosnan, M. et al. Evidence Accumulation Rate Moderates the Relationship between Enriched Environment Exposure and Age-Related Response Speed Declines. Journal of Neuroscience 43, 6401–6414 (2023).

21. Rommelse, N., Luman, M. & Kievit, R. Slow processing speed: a cross-disorder phenomenon with significant clinical value, and in need of further methodological scrutiny. Eur Child Adolesc Psychiatry 29, 1325–1327 (2020).

22. de la Piedra Walter, M., Notbohm, A., Eling, P. & Hildebrandt, H. Audiospatial evoked potentials for the assessment of spatial attention deficits in patients with severe cerebrovascular accidents. J Clin Exp Neuropsychol 43, 623–636 (2021).

23. Di Russo, F., Bozzacchi, C., Matano, A. & Spinelli, D. Hemispheric differences in VEPs to lateralised stimuli are a marker of recovery from neglect. Cortex 49, 931–939 (2013).

24. Di Gregorio, F. et al. Hierarchical psychophysiological pathways subtend perceptual asymmetries in Neglect. Neuroimage 270, 119942 (2023).

25. Lasaponara, S. et al. EEG correlates of preparatory orienting, contextual updating and inhibition of sensory processing in left spatial neglect. Journal of Neuroscience 38, 3792–3808 (2018).

26. Tarkka, I. M., Luukkainen-Markkula, R., Pitkänen, K. & Hämäläinen, H. Alterations in visual and auditory processing in hemispatial neglect: An evoked potential follow-up study. International Journal of Psychophysiology 79, 272–279 (2011).

27. Choinski, M., Szelag, E., Wolak, T. & Szymaszek, A. Neuropsychological correlates of P300 parameters in individuals with aphasia. Int J Lang Commun Disord 58, 256–269 (2023).

28. Doricchi, F. et al. Deficits of hierarchical predictive coding in left spatial neglect. Brain Commun 3, (2021).

29. Gummow, L. J., Dustman, R. E. & Keaney, R. P. Cerebrovascular accident alters P300 event-related potential characteristics. Electroencephalogr Clin Neurophysiol 63, 128–137 (1986).

30. Ladurner, G., Schimke, H., Wranek, U. & Klimesch, W. The value of P300 in the diagnosis of cognitive impairment in stroke. Arch Gerontol Geriatr 10, 1–8 (1990).

31. Priftis, K. et al. Lost in number space after right brain damage: A neural signature of representational neglect. Cortex 44, 449–453 (2008).

32. Wessel, J. R., Klein, T. A., Ott, D. V. M. & Ullsperger, M. Lesions to the prefrontal performance-monitoring network disrupt neural processing and adaptive behaviors after both errors and novelty. Cortex 50, 45–54 (2014).

33. Lasaponara, S., Pinto, M., Aiello, M., Tomaiuolo, F. & Doricchi, F. The hemispheric distribution of α-band eeg activity during orienting of attention in patients with reduced awareness of the left side of space (Spatial neglect). Journal of Neuroscience 39, 4332–4343 (2019).

34. Lasaponara, S. et al. Pre-motor deficits in left spatial neglect: An EEG study on Contingent Negative Variation (CNV) and response-related beta oscillatory activity. Neuropsychologia 147, (2020).

35. Colson, C., Demeurisse, G., Hublet, C. & Slachmuylder, J. L. Subcortical Neglect as a Consequence of a Remote Parieto-Temporal Dysfunction. A Quantitative Eeg Study. Cortex 37, 619–625 (2001).

36. Lhermitte, F., Turell, E., Lebrigand, D. & Chain, F. Unilateral Visual Neglect and Wave P 300: A Study of Nine Cases With Unilateral Lesions of the Parietal Lobes. Arch Neurol 42, 567–573 (1985).

37. Saevarsson, S., Kristjánsson, Á., Bach, M. & Heinrich, S. P. P300 in neglect. Clin Neurophysiol 123, 496–506 (2012).

38. Verleger, R., Heide, W., Butt, C., Wascher, E. & Kömpf, D. On-line brain potential correlates of right parietal patients’ attentional deficit. Electroencephalogr Clin Neurophysiol 99, 444–457 (1996).

39. Rastelli, F. et al. Neural dynamics of neglected targets in patients with right hemisphere damage. Cortex 49, 1989–1996 (2013).

40. Ratcliff, R., Philiastides, M. G. & Sajda, P. Quality of evidence for perceptual decision making is indexed by trial-to-trial variability of the EEG. Proceedings of the National Academy of Sciences 106, 6539–6544 (2009).

41. Britten, K. H., Shadlen, M. N., Newsome, W. T. & Movshon, J. A. The analysis of visual motion: a comparison of neuronal and psychophysical performance. Journal of Neuroscience 12, 4745–4765 (1992).

42. Kelly, S. P. & O’Connell, R. G. Internal and External Influences on the Rate of Sensory Evidence Accumulation in the Human Brain. Journal of Neuroscience 33, 19434–19441 (2013).

43. O’Connell, R. G., Dockree, P. M. & Kelly, S. P. A supramodal accumulation-to-bound signal that determines perceptual decisions in humans. Nature Neuroscience 2012 15:12 15, 1729–1735 (2012).

44. Newman, D. P., Loughnane, G. M., Kelly, S. P., O’Connell, R. G. & Bellgrove, M. A. Visuospatial Asymmetries Arise from Differences in the Onset Time of Perceptual Evidence Accumulation. The Journal of Neuroscience 37, 3378 (2017).

45. Twomey, D. M., Murphy, P. R., Kelly, S. P. & O’Connell, R. G. The classic P300 encodes a build-to-threshold decision variable. Eur J Neurosci 42, 1636–1643 (2015).

46. Kelly, S. P., Corbett, E. A. & O’connell, R. G. Neurocomputational mechanisms of prior-informed perceptual decision-making in humans. Nat Hum Behav (2021) doi:10.1038/s41562-020-00967-9.

47. Murphy, P. R., Robertson, I. H., Harty, S. & O’Connell, R. G. Neural evidence accumulation persists after choice to inform metacognitive judgments. Elife 4, e11946 (2015).

48. Steinemann, N. A., O’Connell, R. G. & Kelly, S. P. Decisions are expedited through multiple neural adjustments spanning the sensorimotor hierarchy. Nat Commun 9, (2018).

49. Twomey, D. M., Kelly, S. P. & O’Connell, R. G. Abstract and Effector-Selective Decision Signals Exhibit Qualitatively Distinct Dynamics before Delayed Perceptual Reports. J Neurosci 36, 7346–7352 (2016).

50. Ruesseler, M., Weber, L. A., Marshall, T. R., O’Reilly, J. & Hunt, L. T. Quantifying decision-making in dynamic, continuously evolving environments. Elife 12, (2023).

51. Herding, J., Ludwig, S., von Lautz, A., Spitzer, B. & Blankenburg, F. Centro-parietal EEG potentials index subjective evidence and confidence during perceptual decision making. Neuroimage 201, 116011 (2019).

52. van Vugt, M. K., Beulen, M. A. & Taatgen, N. A. Relation between centro-parietal positivity and diffusion model parameters in both perceptual and memory-based decision making. Brain Res 1715, 1–12 (2019).

53. von Lautz, A., Herding, J. & Blankenburg, F. Neuronal signatures of a random-dot motion comparison task. Neuroimage 193, 57–66 (2019).

54. Spitzer, B., Waschke, L. & Summerfield, C. Selective overweighting of larger magnitudes during noisy numerical comparison. Nat Hum Behav 1, 1–8 (2017).

55. Philiastides, M. G., Heekeren, H. R. & Sajda, P. Human Scalp Potentials Reflect a Mixture of Decision-Related Signals during Perceptual Choices. The Journal of Neuroscience 34, 16877 (2014).

56. McGovern, D. P., Hayes, A., Kelly, S. P. & O’Connell, R. G. Reconciling age-related changes in behavioural and neural indices of human perceptual decision-making. Nat Hum Behav 2, 955– 966 (2018).

57. Becker, E. & Karnath, H. O. Incidence of Visual Extinction After Left Versus Right Hemisphere Stroke. Stroke 38, 3172–3174 (2007).

58. Stefanac, N. R., Zhou, S. H., Spencer-Smith, M. M., O’Connell, R. & Bellgrove, M. A. A neural index of inefficient evidence accumulation in dyslexia underlying slow perceptual decision making. Cortex 142, 122–137 (2021).

59. Van Kempen, J. et al. Behavioural and neural signatures of perceptual decision-making are modulated by pupil-linked arousal. Elife 8, (2019).

60. Prigatano, G. P. Neuropsychological rehabilitation after brain injury. (No Title) (Johns Hopkins University Press, 1986).

61. Barskova, T. & Wilz, G. Psychosocial functioning after stroke: Psychometric properties of the patient competency rating scale. Brain Inj 20, 1431–1437 (2006).

62. Corbetta, M. & Shulman, G. L. Control of goal-directed and stimulus-driven attention in the brain. Nature Reviews Neuroscience 2002 3:3 3, 201–215 (2002).

63. Shadlen, M. N. & Kiani, R. Decision Making as a Window on Cognition. Neuron 80, 791–806 (2013).

64. Jørgensen, H. S. et al. Outcome and time course of recovery in stroke. Part II: Time course of recovery. The copenhagen stroke study. Arch Phys Med Rehabil 76, 406–412 (1995).

65. Xi, G., Keep, R. F. & Hoff, J. T. Mechanisms of brain injury after intracerebral haemorrhage. Lancet Neurol 5, 53–63 (2006).

66. Hankey, G. J. Stroke. The Lancet 389, 641–654 (2017).

67. Bernhardt, J. et al. Agreed Definitions and a Shared Vision for New Standards in Stroke Recovery Research: The Stroke Recovery and Rehabilitation Roundtable Taskforce. Neurorehabil Neural Repair 31, 793–799 (2017).

68. Lindell, A. B. et al. Clinical assessment of hemispatial neglect: evaluation of different measures and dimensions. Clin Neuropsychol 21, 479–497 (2007).

69. Gherman, S. et al. Intracranial electroencephalography reveals effector-independent evidence accumulation dynamics in multiple human brain regions. Preprint at 10.1101/2023.04.10.536314.

70. Price, C. J., Hope, T. M. & Seghier, M. L. Ten problems and solutions when predicting individual outcome from lesion site after stroke. Neuroimage 145, 200 (2017).

71. Nasreddine, Z. S., et al. The Montreal Cognitive Assessment, MoCA: A Brief Screening Tool For Mild Cognitive Impairment. J Am Geriatr Soc 53, 695–699 (2005).

72. Demeyere, N., Riddoch, M. J., Slavkova, E. D., Bickerton, W. L. & Humphreys, G. W. The Oxford Cognitive Screen (OCS): validation of a stroke-specific short cognitive screening tool. Psychol Assess 27, 883–894 (2015).

73. Bender, M. B. Disorders in Perception: With Particular Reference to the Phenomena of Extinction and Displacement. Disorders in perception: With particular reference to the phenomena of extinction and displacement. (Charles C Thomas Publisher, 2011). doi:10.1037/13218-000.

74. Nelson, H. E. & Willison, J. National Adult Reading Test (NART). (1991).

75. Wechsler, D. Wechsler Adult Intelligence Scale, Fourth Edition (WAIS-IV). (2008).

76. Bright, P., Hale, E., Gooch, V. J., Myhill, T. & van der Linde, I. The National Adult Reading Test: restandardisation against the Wechsler Adult Intelligence Scale—Fourth edition. Neuropsychol Rehabil 28, 1019–1027 (2016).

77. Nucci, M., Mapelli, D. & Mondini, S. Cognitive Reserve Index questionnaire (CRIq): a new instrument for measuring cognitive reserve. Aging Clin Exp Res 24, 218–226 (2012).

78. Brainard, D. H. The Psychophysics Toolbox. Spat Vis 10, 433–436 (1997).

79. Cornelissen, F. W., Peters, E. M. & Palmer, J. The Eyelink Toolbox: Eye tracking with MATLAB and the Psychophysics Toolbox. Behavior Research Methods, Instruments, and Computers 34, 613–617 (2002).

80. Pelli, D. G. The VideoToolbox software for visual psychophysics: Transforming numbers into movies. Spat Vis 10, 437–442 (1997).

81. Humphreys, G., Bickerton, W.-L., Samson, D. & Riddoch, M. BCoS Cognition Screen. (2012).

82. Bellgrove, M. A. et al. Association between Dopamine Transporter (DAT1) Genotype, Left-Sided Inattention, and an Enhanced Response to Methylphenidate in Attention-Deficit Hyperactivity Disorder. Neuropsychopharmacology 30, 2290–2297 (2005).

83. Newsome, W. T., Britten, K. H. & Movshon, J. A. Neuronal correlates of a perceptual decision. Nature 341, 52–54 (1989).

84. Kosgallana, A., Cordato, D., Kam, D., Chan, Y. & Yong, J. Use of Cognitive Screening Tools to Detect Cognitive Impairment After an Ischaemic Stroke: a Systematic Review. SN Comprehensive Clinical Medicine 2018 1:4 1, 255–262 (2019).

85. Moore, M. et al. Rapid screening for neglect following stroke: A systematic search and European Academy of Neurology recommendations. Eur J Neurol 29, 2596–2606 (2022).

86. Bickerton, W. L. et al. The BCoS cognitive profile screen: Utility and predictive value for stroke. Neuropsychology 29, 638–648 (2015).

87. Moore, M. J. et al. A comparison of lesion mapping analyses based on CT versus MR imaging in stroke. Neuropsychologia 184, 108564 (2023).

88. Moore, M. J. & Demeyere, N. Lesion symptom mapping of domain-specific cognitive impairments using routine imaging in stroke. Neuropsychologia 167, (2022).

89. Moore, M. J. A Practical Guide to Lesion Symptom Mapping. doi:10.31234/OSF.IO/2JXR9.

90. Rorden, C., Karnath, H. O. & Bonilha, L. Improving lesion-symptom mapping. J Cogn Neurosci 19, 1081–1088 (2007).

91. Penny, W., Friston, K., Ashburner, J., Kiebel, S. & Nichols, T. Statistical Parametric Mapping: The Analysis of Functional Brain Images. Statistical Parametric Mapping: The Analysis of Functional Brain Images (2007) doi:10.1016/B978-0-12-372560-8.X5000-1.

92. Rorden, C., Bonilha, L., Fridriksson, J., Bender, B. & Karnath, H. O. Age-specific CT and MRI templates for spatial normalization. Neuroimage 61, 957–965 (2012).

93. Moore, M. J., Milosevich, E., Mattingley, J. B. & Demeyere, N. The neuroanatomy of visuospatial neglect: A systematic review and analysis of lesion-mapping methodology. Neuropsychologia 180, 108470 (2023).

94. Schaefer, A. et al. Local-Global Parcellation of the Human Cerebral Cortex from Intrinsic Functional Connectivity MRI. Cereb Cortex 28, 3095–3114 (2018).

95. Griffis, J. C., Metcalf, N. V., Corbetta, M. & Shulman, G. L. Lesion Quantification Toolkit: A MATLAB software tool for estimating grey matter damage and white matter disconnections in patients with focal brain lesions. Neuroimage Clin 30, 102639 (2021).

96. Tzourio-Mazoyer, N. et al. Automated Anatomical Labeling of Activations in SPM Using a Macroscopic Anatomical Parcellation of the MNI MRI Single-Subject Brain. Neuroimage 15, 273–289 (2002).

97. Delorme, A. & Makeig, S. EEGLAB: An open source toolbox for analysis of single-trial EEG dynamics including independent component analysis. J Neurosci Methods 134, 9–21 (2004).

98. Kayser, J. & Tenke, C. E. Principal components analysis of Laplacian waveforms as a generic method for identifying ERP generator patterns: II. Adequacy of low-density estimates. Clinical Neurophysiology 117, 369–380 (2006).

