## Supplemental Materials for "Neurophysiological mechanisms underlying post-stroke deficits in contralesional perceptual processing"

**Supplementary Materials**

**Supplementary Figure 1**

*Normalised Lesion Masks for Individual Patients*

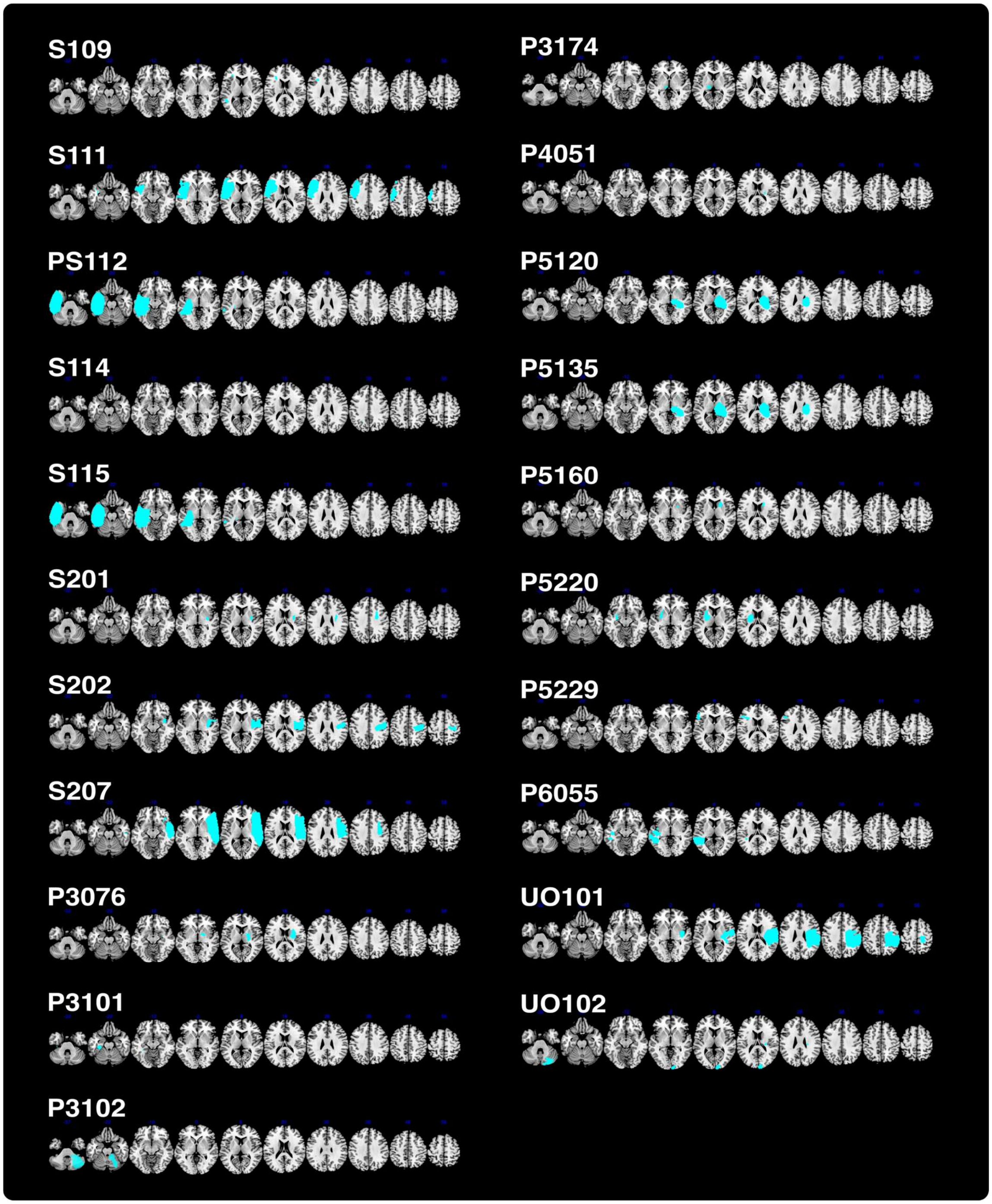

**Supplementary Figure 2**

*Network Disconnection Summaries for Individual Patients*

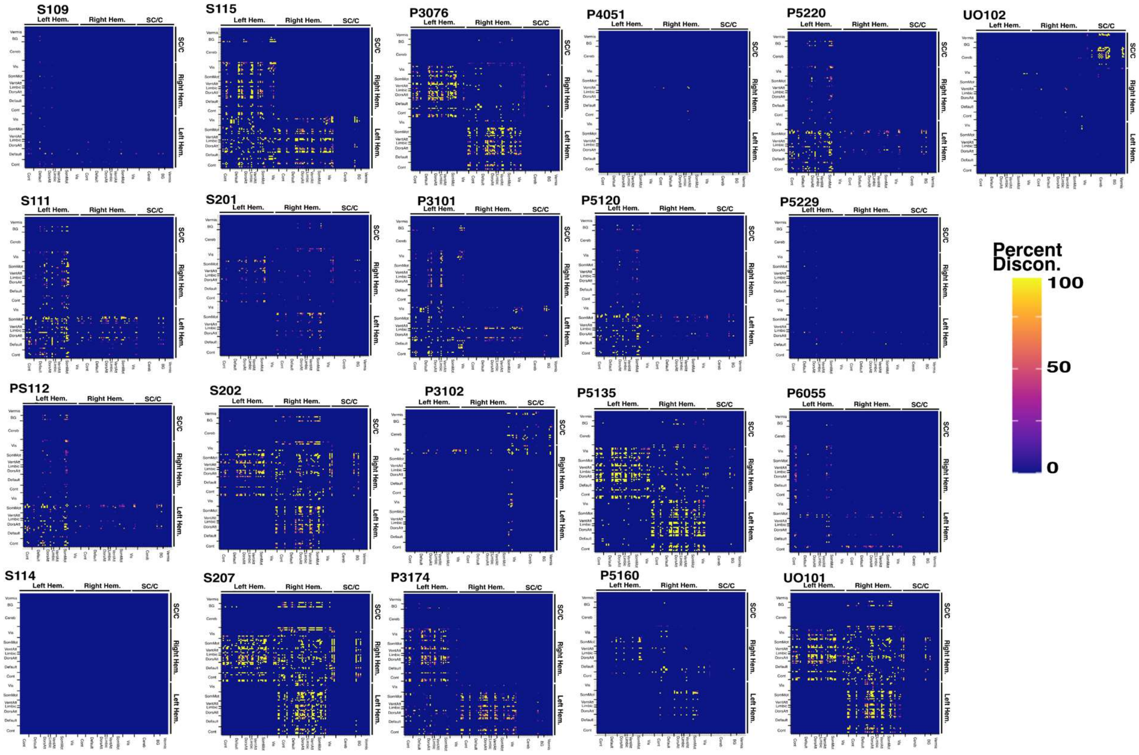


**Supplementary Figure 3**

*Relationships Between Self-Reported Everyday Functioning and Neurophysiological Markers*

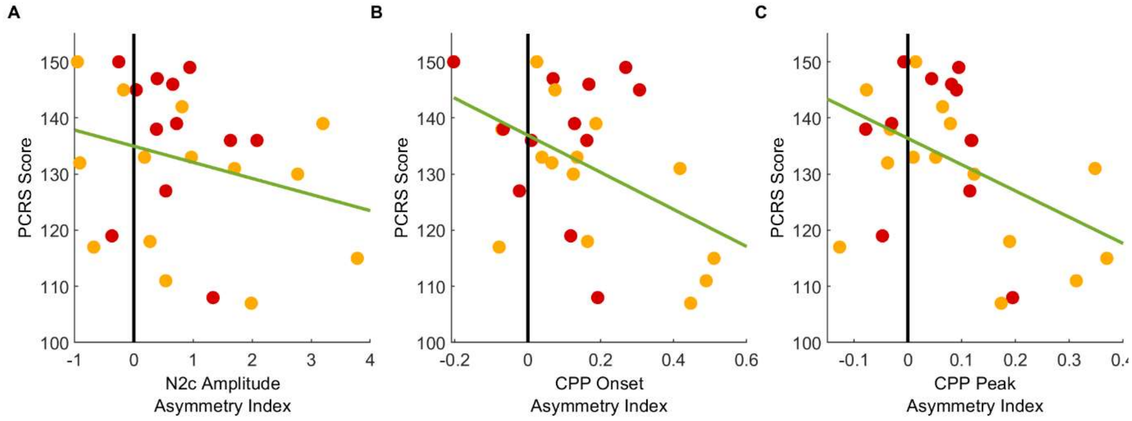


Scatterplots depict the relationship between self-reported levels of everyday functioning on the PCRS and individual subject-level mean asymmetries for (**A**) N2c amplitude, (**B**) CPP onset, and (**C**) CPP peak latency. A least-squares regression line is fitted for reference.

### Supplementary Tables

#### Supplementary Table 1

*Results of Linear Mixed Models for Group and Target Side Across Measures of RT, N2c Amplitude, and CPP Peak Amplitude*

| **Model: RT ~ 1 + Hemifield*Group + (1 + Hemifield\|Subject)** | | | | |
|---|---|---|---|---|
| **Parameter** | **Fixed Effects Estimate (*SE*)** | **95% CI** | **Test (*df*)** | ***p*** |
| Group (reference = healthy older) | | | *F*(2,11275)=3.87 | .02 |
| Left Stroke | -6.69 (*28.96*) | -63.46, 50.07 | *t*(11275)=-0.23 | .82 |
| Right Stroke | -78.69 (*35.27*) | -147.83, -9.55 | *t*(11275)=-2.23 | .03 |
| Hemifield (reference = left) | | | *F*(1,11275)=2.27 | .13 |
| Right Hemifield | 14.90 (*9.89*) | -4.49, 34.29 | *t*(11275)=1.51 | .13 |
| Group x Hemifield | | | *F*(2,11275)=7.21 | <.001 |
| Left Stroke | -15.09 (*12.64*) | -39.87, 9.68 | *t*(11275)=-1.19 | .23 |
| Right Stroke | -37.34 (*15.38*) | -67.49, -7.19 | *t*(11275)=-2.43 | .02 |
| **Random Effects** | | | | |
| **Parameter** | **Variance** | | **SD** | |
| Intercept | 26116.16 | | 214.64 | |
| Hemifield | 4827.84 | | 138.97 | |
| **Model Fit ($R^2$)** | | | | |
| **Ordinary** | | **Adjusted** | | |
| 0.42 | | 0.41 | | |

| **Model: N2c Amplitude ~ 1 + Group*Hemifield + (1 + Hemifield\|Subject)** | | | | |
|---|---|---|---|---|
| **Parameter** | **Fixed Effects Estimate (*SE*)** | **95% CI** | **Test (*df*)** | ***p*** |
| Group (reference = healthy older) | | | *F*(2,8599)=1.35 | .26 |
| Left Stroke | -2.52 (*1.73*) | -5.92, 0.87 | *t*(8599)=-1.46 | .15 |
| Right Stroke | 0.29 (*2.11*) | -3.85, 4.43 | *t*(8599)=0.13 | .89 |
| Hemifield (reference = left) | | | *F*(1,8599)=0.00005 | .99 |

| | | | | |
|---|---|---|---|---|
| Right Hemifield | 0.007 (*1.02*) | -2.00, 2.02 | *t*(8599)=0.007 | .99 |
| Group x Hemifield | | | *F*(2,8599)=6.25 | .002 |
| Left Stroke | -1.03 (*1.31*) | -3.60, 1.53 | *t*(8599)=-0.79 | .43 |
| Right Stroke | -3.98 (*1.60*) | -7.11, -0.85 | *t*(8599)=-2.49 | .01 |
| **Random Effects** | | | | |
| **Parameter** | **Variance** | | **SD** | |
| Intercept | 87.14 | | 9.33 | |
| Hemifield | 47.08 | | 6.86 | |
| **Model Fit ($R^2$)** | | | | |
| **Ordinary** | **Adjusted** | | | |
| 0.14 | 0.14 | | | |

**Model: CPP Peak Amplitude ~ Group*Hemifield + (1|Subject)**

| **Parameter** | **Fixed Effects**<br>**Estimate (*SE*)** | **95% CI** | **Test (*df*)** | ***p*** |
|---|---|---|---|---|
| Group (reference = healthy older) | | | *F*(2,11091)=2.42 | .09 |
| Left Stroke | -4.11 (*1.95*) | -7.93, -0.28 | *t*(11091)=-2.10 | .04 |
| Right Stroke | 3.94 (*2.37*) | -0.70, 8.59 | *t*(11091)=1.66 | .10 |
| Hemifield (reference = left) | | | *F*(1,11091)=2.85 | .09 |
| Right Hemifield | 1.04 (*0.61*) | -0.17, 2.24 | *t*(11091)=1.69 | .09 |
| Group x Hemifield | | | *F*(2,11091)=2.45 | .09 |
| Left Stroke | -0.08 (*0.79*) | -1.64, 1.47 | *t*(11091)=-0.11 | .92 |
| Right Stroke | 1.84 (*0.95*) | -0.02, 3.70 | *t*(11091)=1.93 | .05 |
| **Random Effects** | | | | |
| **Parameter** | **Variance** | | **SD** | |
| Intercept | 103.73 | | 10.19 | |
| Hemifield | 4.23 | | 2.06 | |
| **Model Fit ($R^2$)** | | | | |
| **Marginal** | **Conditional** | | | |
| 0.03 | 0.03 | | | |

**Supplementary Table 2**

*Regression Coefficients for Each Predictor Variable on BRDM RT, PCRS Score, and Asymmetry Indices for RT, N2c Amplitude, and CPP Onset and Peak Latency*

| **Predictor Variable** | ***B* [95% CI]** | **$R^2$ (adjusted)** |
|---|---|---|

| | RT Asymmetry (ms) | |
|---|---|---|
| **N2c Amplitude Asymmetry** | 0.09 [0.04, 0.15] | 0.47 (0.43) |
| **Group** | 0.07 [-0.09, 0.22] | |
| **N2i Amplitude Asymmetry** | -0.003 [-0.04, 0.04] | 0.24 (0.19) |
| **Group** | 0.21 [0.05, 0.37] | |
| **CPP Onset Latency Asymmetry** | 0.81 [0.49, 1.13] | 0.64 (0.61) |
| **Group** | 0.001 [-0.14, 0.14] | |
| **CPP Peak Latency Asymmetry** | 1.36 [1.04, 1.68] | 0.80 (0.79) |
| **Group** | 0.01 [-0.09, 0.10] | |
| | **PCRS** | |
| **RT Asymmetry** | -27.71 [-51.74, -3.69] | 0.25 (0.19) |
| **Group** | -5.22 [-14.88, 4.45] | |
| **N2c Amplitude Asymmetry** | -2.57 [-6.95, 1.81] | 0.13 (0.06) |
| **Group** | -6.63 [-16.93, 3.68] | |
| **CPP Onset Latency Asymmetry** | -29.17 [-57.72, -0.61] | 0.24 (0.18) |
| **Group** | -5.64 [-15.79, 4.51] | |
| **CPP Peak Latency Asymmetry** | -43.27 [-81.99, -4.56] | 0.24 (0.18) |
| **Group** | -5.55 [-15.23, 4.13] | |

### Supplementary Results

#### Supplementary Results 1 – Contralesional Slowing is Present in Those With and Without Neglect

We sought to determine whether our findings of contralesional slowing were driven by the subset of right hemisphere stroke participants who had clinically significant neglect, as evidenced by performance on a cancellation task ($n$=7), as compared with those who did not show evidence of neglect ($n$=11). We used a two-way mixed ANOVA to investigate the effects of group (Neglect, No Neglect) and target hemifield (Left, Right) on RT. There was no significant Group x Hemifield interaction, $F(1,16)=0.002$, $p=.96$, indicating that right hemisphere patients both with and without clinically apparent neglect respond more slowly to stimuli presented in the contralesional (left) hemifield.

**Supplementary Results 2 – Marginal Evidence for Relationships Between Evidence Accumulation Asymmetries and Post-Stroke Everyday Functioning**

To examine whether our neurophysiological asymmetry markers were associated with self-reported everyday functioning, we first re-calculated CPP onset latency and CPP peak latency asymmetries in accordance with our process for RT. This transformation means that greater CPP onset latency and CPP peak latency asymmetries indicate slower responses to contralesional stimuli, regardless of lesion side. We then ran three separate multiple regressions with PCRS score as the outcome variable, the above asymmetry measures as predictors, and stroke side as a covariate. Note that one participant had an invalid CPP onset, hence different degrees of freedom cited below.

There was no significant relationship between N2c amplitude asymmetry and everyday functioning, $F(1,24)=1.47$, $p_{corr}=.24$. Once correcting for multiple comparisons using Bonferroni-Holm-adjusted $p$ values, there was no significant relationship between everyday functioning and CPP onset asymmetry ($F(1,23)=4.47$, $p_{corr}=.09$) or CPP peak asymmetry ($F(1,24)=5.32$, $p_{corr}=.09$; Supplementary Fig. 1).
